# ChromSMF: integrated profiling of histone modifications, protein-DNA interactions and DNA methylation on multi-kilobase DNA molecules

**DOI:** 10.64898/2026.03.11.710921

**Authors:** Michela Palamin, Arnaud R. Krebs

**Affiliations:** Genome Biology Unit, EMBL Heidelberg, Meyerhofstraße 1, 69117 Heidelberg, Germany; Faculty of Biosciences, Collaboration for Joint PhD Degree between EMBL and Heidelberg University, Heidelberg, Germany

## Abstract

Chromatin accessibility and histone post-translational modifications are widely used to identify and characterize *cis*-regulatory elements. Yet, these are typically measured separately, precluding direct linkage between them. Here, we present Chromatin-informed Single Molecule Footprinting (ChromSMF), a method that simultaneously quantifies histone modifications, transcription factor binding, and nucleosome occupancy, while measuring DNA methylation and sequence variations on the same multi-kilobase DNA molecules. ChromSMF combines antibody-guided tethering of the adenine methyltransferase Hia5 to modified histones, protein-DNA footprinting using the cytosine methyltransferase M.CviPI, and direct methylation detection by Nanopore sequencing. We benchmark ChromSMF across five histone modifications, uncovering relationships between epigenetic states and chromatin opening across entire *cis*-regulatory landscapes. We further present a computational framework for integrated analysis of multiple layers of epigenetic regulation on individual haplotypes. Together, ChromSMF provides an integrated genome-wide genomic method to investigate the combinatorial function of multiple genetic and epigenetic factors on gene regulation across diverse cellular contexts.

## Introduction

Epigenetic marks, such as DNA methylation and histone modifications are essential regulators of gene expression during development, ensuring proper cell differentiation and tissue formation (1, 2). Perturbations of the deposition, removal, or recognition of these marks are associated with numerous pathologies. For instance, histone-modifying enzymes are frequently mutated in cancer (3, 4), and specific alterations in the DNA methylation landscape are characteristic of aging (5). Chemical and CRISPR-based epigenome editing have recently emerged as an exciting opportunity to reverse disease-associated changes in histone modification distribution (6). However, for such approaches to succeed and restore physiological transcriptional programs, the targeted histone modification must functionally modulate downstream regulatory processes, such as chromatin opening and transcription factor (TF) binding at *cis*-regulatory elements (cREs). The regulatory effects of individual histone modifications not only vary between their type, but it is also influenced by the genomic context where they occur (*i*.*e*., enhancers and promoters) (1, 7–9). In addition, genetic variation between individuals can further influence epigenetic states, contributing to differences in gene regulation and disease susceptibility (10–12). Therefore, elucidating how distinct histone modifications regulate cRE activity in specific genomic contexts remains a fundamental question, necessitating approaches that directly associate the presence or absence of specific histone modifications with chromatin accessibility, TF occupancy, and variation in the underlying genetic sequence.

Chromatin accessibility and histone modifications are generally quantified through the enrichment of short DNA fragments corresponding to their associated loci. For instance, ChIP-seq, CUT&RUN, and derived methods, efficiently map histone modifications or TFs (13–15). Similarly, while ATAC-seq and DNase-seq are widely used to profile chromatin accessibility genome-wide (16, 17), MNaseseq maps nucleosome positioning at near base-pair resolution (18, 19). These methods have revealed that the presence of histone modifications tightly correlates with the chromatin accessibility of cREs as well as with the activity of the regulated genes. For instance, H3K4me1 and H3K4me3 associate with the activation of enhancers and promoters, respectively (20, 21). However, these associations remain correlative as the presence or absence of a particular histone modification and the resulting changes in chromatin accessibility and TF binding cannot be resolved simultaneously in the same assay.

Single-molecule genomic methods have recently emerged as powerful approaches to overcome these limitations, as they enable simultaneous measurement of multiple regulatory features on individual DNA molecules (22). The frequency of co-occurrence of two genomic events on the same molecules can then be used to infer positive, negative, or neutral associations between them (8, 22–26); an information that is lost when correlating bulk experiments performed individually. For example, Single Molecule Footprinting (SMF), which employs exogenous cytosine methyltransferases to map the simultaneous occupancy of multiple TFs across the genome, identified cooperativity between them (24). By simultaneously measuring DNA methylation and chromatin accessibility on the same DNA molecules, SMF further identified the subset of enhancers that are epigenetically repressed by DNA methylation (8). Footprinting technologies combining methyltransferases (cytosine, or adenine), or deaminases, with long-read sequencing have extended the analysis of molecular co-occurrence to loci separated by several kilobases (23, 27–39). In parallel, assays based on antibody-directed adenine methylation, such as DiMeLo-seq, have enabled targeted detection of specific histone modifications and DNA-binding factors at single-molecule resolution (40–44). Despite these advances, current approaches remain constrained in the number of molecular layers that can be simultaneously profiled, and computational frameworks to quantitatively assess their molecular co-occurrence are lacking. Consequently, systematically establishing the functional consequences of histone modification deposition on other regulatory layers, such as chromatin opening or TF binding, as well as their association with underlying genetic variation, has remained challenging.

To address this gap, we developed Chromatin-informed Single Molecule Footprinting (ChromSMF), an integrated long-read single-molecule genomic approach that simultaneously detects 1) histone modifications, 2) chromatin accessibility, 3) TF binding, 4) endogenous DNA methylation, and 5) genetic variation on the same, individual DNA molecules genome-wide. ChromSMF combines mapping of histone modifications through antibody-directed adenine methylation (mA), DNA footprinting of nucleosomes and TFs via methylation of accessible cytosines (mC), and direct methylation profiling using Nanopore sequencing. Here, we demonstrate ChromSMF ability to profile three active (H3K4me3, H3K27ac, H3K4me1) and two repressive (H3K27me3, H3K9me3) histone modifications across distinct cellular contexts. We establish a computational framework that integrates these five molecular layers at singlemolecule resolution, enabling quantitative analysis of their combinatorial relationships. We illustrate the power of this approach in defining the specific chromatin accessibility and nucleosome occupancy associated with individual histone modifications across entire *cis*-regulatory landscapes. Finally, we extend ChromSMF to haplotype-resolved analysis in mammalian genomes, and demonstrate its utility in quantitatively comparing genetic and epigenetic mechanisms to assess their relative contributions to allelic differences in gene expression.

## Results

### A tandem assay to simultaneously measure histone modifications and chromatin accessibility

ChromSMF involves the sequential treatment of permeabilized cells with the exogenous GpC methyltransferase (MTase) M.CviPI, followed by the antibody-directed adenine MTase Hia5 (Figure 1A). An additional CpG MTase (M.SssI) treatment can optionally be included to increase the spatial resolution of the footprinting (from 14 to 7 bp (45)) in genomes lacking endogenous methylation, such as *Drosophila*. First, cells are immobilized on Concanavalin A beads and chemically permeabilized. Then, incubation with cytosine MTases pervasively deposits methylation at accessible chromatin regions, whereas DNA protected by histones or TFs remains unmethylated (adapted from SMF (45, 46)). Second, sequential incubations with a primary antibody against the target of interest (*e*.*g*., a histone modification, HM) and the pA-Hia5 fusion protein result in localised adenine methylation at loci harbouring the targeted HM (similar to DiMeLo-seq (40, 42)). To minimize background methylation, MTases activation is controlled through multiple rounds of addition and washing of their obligatory co-factor S-adenosylmethionine (SAM). Finally, the resulting footprinted genomic DNA is extracted and prepared for direct methylation profiling using Oxford Nanopore Technology (ONT) sequencing, that simultaneously detects cytosine (mC) and adenine (mA) methylation on the same DNA molecules (Figure 1A). The resulting methylation can then be analysed at bulk level by averaging methylation at individual cytosines and adenines to detect chromatin accessibility and HM, respectively (Figure 1A; right upper panel). Additionally, integration of the two signals from the same individual DNA molecules allows the identification of the fraction of molecules at which multiple regulatory layers co-occur in a cell population (Figure 1A; right lower panel).

**Fig. 1.**
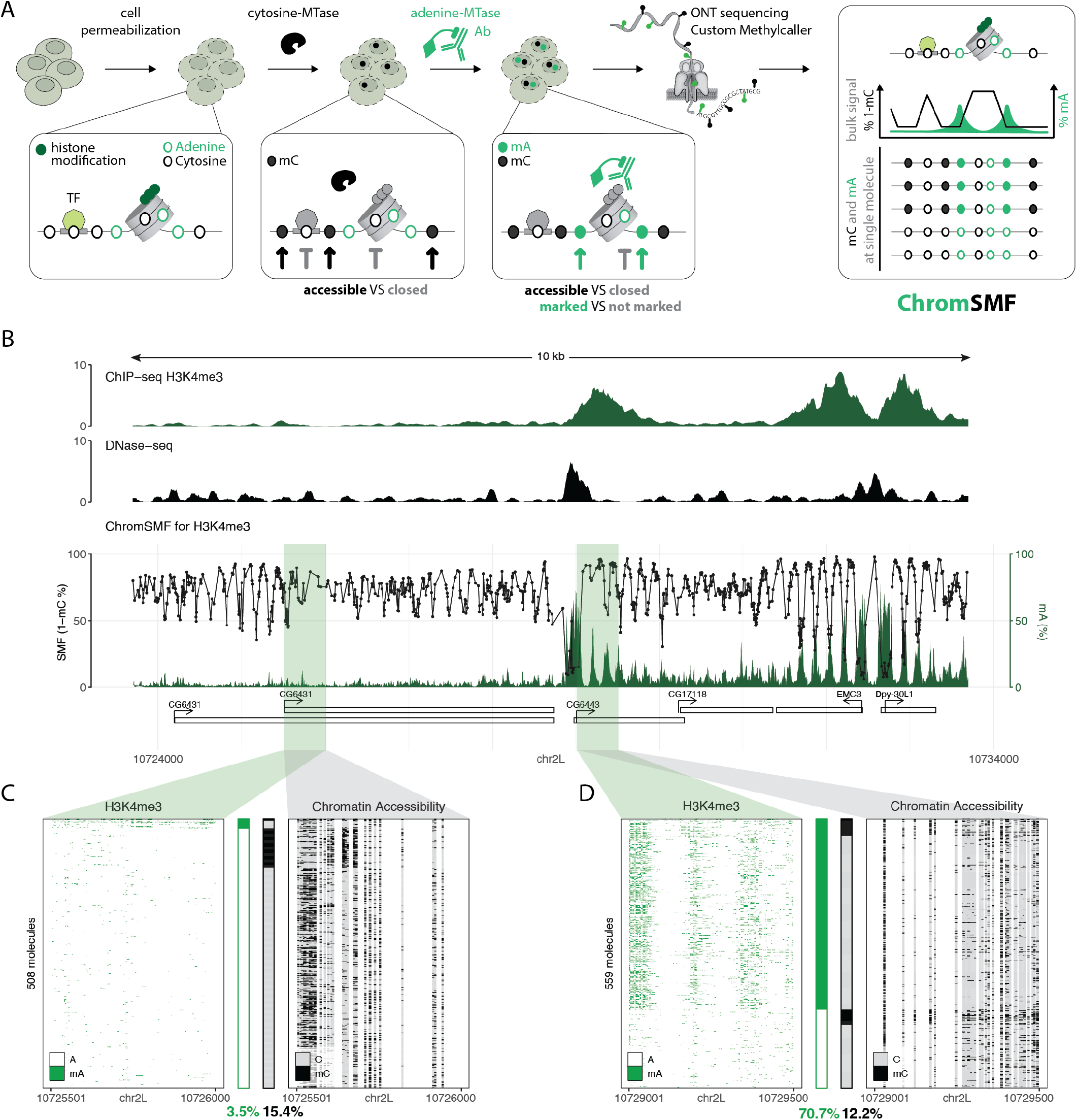
ChromSMF simultaneously measures histone modifications and chromatin accessibility in a single assay. **(A) Schematic representation of the two-step ChromSMF protocol**. Cells are chemically permeabilized and incubated with M.CviPI (cytosine-MTase), a methyltransferase that deposits cytosine methylation at accessible GpCs across the genome (black spheres). The obligatory cofactor SAM is washed out to stop M.CviPI activity. Cells are then sequentially incubated with an antibody (Ab) targeting a histone modification, and the proteinA-Hia5 (adenine-MTase) fusion protein that will methylate adenines in proximity of the histone modification of interest (green spheres). Sequencing of the resulting DNA using Oxford Nanopore Technologies (ONT) enables simultaneous detection of histone modifications (green signal; mA) and transcription factor binding events (black signal; mC) at bulk and single-molecule resolution. **(B) Example locus illustrating the simultaneous detection of H3K4me3 and chromatin accessibility at active promoters**. Top panel: genome browser tracks displaying ChIP-seq enrichment for H3K4me3 (green) and DNase-seq signal (black). Lower panel: ChromSMF sample for H3K4me3. Average SMF signal (1 – mC%) of individual cytosines (black) and average smoothed mA signal (mA%; green; smoothing across 4 adenines). **(C) Simultaneous detection of chromatin accessibility and H3K4me3 on individual DNA molecules at a locus with low ChIP-seq enrichment for H3K4me3**. Single-molecule stacks display either mA-H3K4me3 (green, left) or mC-chromatin accessibility (black, right) signal. Molecules are displayed in identical order in both panels and originate from the same sample. Single-molecule classification of H3K4me3 (green) and chromatin accessibility (black) are shown as stacked bar plots between the single-molecule stacks. Single-molecule quantification of total H3K4me3 and chromatin accessibility at the locus are shown at the bottom. **(D) Simultaneous detection of chromatin accessibility and H3K4me3 on individual DNA molecules at a locus with high ChIP-seq enrichment for H3K4me3**. Single-molecule stacks display either mA-H3K4me3 (green, left) or mC-chromatin accessibility (black, right) signal. Molecules are displayed in identical order in both panels and originate from the same sample. Single-molecule classification of H3K4me3 (green) and chromatin accessibility (black) are shown as stacked bar plots between the single-molecule stacks. Single-molecule quantification of total H3K4me3 and chromatin accessibility at the locus are shown at the bottom.

Similar to other single-molecule genomic methods (33, 40, 45, 47), ChromSMF does not enrich for DNA molecules associated with specific chromatin features (Figure 1A). Instead, DNA molecules are sequenced irrespective of their mC or mA marking status. Therefore, robust estimation of the abundance of a genomic feature requires to cover each locus with dozens of individual reads (45). To develop and benchmark ChromSMF, we initially targeted H3K4me3 in *Drosophila* Schneider’s 2 (S2) cells. H3K4me3 is a histone modification for which well-validated antibodies are available, and its characteristic localization at gene promoters is known (21). In addition, S2 cells are a well-established model for studying chromatin regulation of transcription (48, 49), and their small genome size (< 200 Mb) eases the iterative generation of datasets with sufficient coverage for robust benchmarking. Indeed, a coverage of about 500x can be easily reached using a single PromethION flow cell (typical output above 100 Gb).

ChromSMF uses mC and mA as independent channels to quantify chromatin accessibility and presence of a HM, respectively. It is thus essential to ensure that there is no cross-interference between their detection. Existing ONT methylation caller models were optimised to detect methylation in individual contexts, and the simultaneous presence of mA and mC has been reported to impair calling accuracy (50). We therefore tested whether, in our setup, the presence of one modification interferes with detection of the other when using these models (23). We *in vitro* methylated naked DNA using mC and mA MTases either individually or in combination (see Methods), sequenced it, and called methylation. The accuracy of the model trained to detect mC alone (23) was not affected by the addition of mA (accuracy=0.84, Figure S1A). In contrast, the accuracy of the mA model (ONT sup v4.2) was significantly reduced in the presence of mC, with a false-negative rate increasing to 36% (accuracy=0.77, Figure S1B). To overcome this issue, we trained a custom model for calling mA in the presence of mC (see Methods). This substantially improved mA detection in presence of mC, and reduced the false-negative rate to less than 13% (accuracy=0.92, Figure S1C).

We next optimized the experimental conditions to combine mC-based footprinting with antibody-directed mA deposition. We verified that neither readout was compromised, for example by enzyme competition, or extended incubation times. To that end, we compared the distribution of per-read cytosine methylation rates and benchmarked it against reference datasets in which chromatin accessibility (CA) was measured individually (23). We first attempted to combine the two experimental steps by simultaneously activating the mC MTase and the antibody-directed mA MTase. This resulted in a strong reduction in mC-based footprinting efficiency (Figure S1D). Reasoning that the mA MTase may compete the mC MTase, we next tested a sequential configuration in which the 15 minutes mC-based footprinting precedes the overnight antibody-directed mA. We added a washing step between the treatments in order to deplete SAM and inactivate the mC MTases, thus preventing the extension of the footprinting reaction. This sequential protocol faithfully recapitulated the per-read mC-based footprinting distribution (Figure S1D) and showed consistent genome-wide average mC in both methylated contexts (R > 0.85, Figure S1E-F). Similarly, the H3K4me3-targeted mA levels in our combined protocol were consistent with single-treatment reference data (R = 0.90, Figure S1G). Finally, mC and mA deposition were highly reproducible between biological replicates, demonstrating the robustness of this sequential ChromSMF protocol (R > 0.95, Figure S1H-I).

To confirm that ChromSMF accurately detects both H3K4me3 and CA, we compared them with orthogonal datasets. Inspection of individual example loci showed antibody-directed mA signal occurs at active promoters enriched for H3K4me3 by ChIP-seq, while inactive promoters show low mA levels (Figure 1B; green tracks in ChIP-seq and ChromSMF). This observation was consistent genomewide as increasing levels of H3K4me3 enrichment correlated with higher levels of mA at promoters (Figure S1J). Similarly, average mC profiles recapitulated CA as measured by DNase-seq, detecting increased accessibility upstream of active promoters, and resolving phased nucleosomes in the body of active genes (Figure 1B; black tracks in DNase-seq and ChromSMF; Figure S1K). Together, these results demonstrate that ChromSMF accurately maps both H3K4me3 and CA with performance comparable to that of individual assays, while unifying these readouts within a single experiment.

### Quantification of histone modifications at single– molecule resolution

Unlike bulk assays, ChromSMF does not enrich for chromatin molecules marked by specific HMs. Instead, its specific mA labelling allows direct measure of the molecular heterogeneity in HM deposition at individual loci (Figure 1C-D; left panel). To leverage this property, we built on our existing pipelines for single-molecule quantification of CA (23), and developed a computational strategy to quantify the deposition of H3K4me3 on single DNA molecules (Figure S2A-B). The mC-based footprinting step of our protocol is designed to reach near-saturation levels (45). Consequently, the resulting chromatin accessibility data can be interpreted at the resolution of individual cytosines (23, 45, 46). In contrast, when inspecting individual molecules at H3K4me3-positive promoters (Figure S2C), we noticed that only a fraction of the adenines was methylated on each molecule (Figure 1D left panel; Figure S2A-B). Nevertheless, mA frequency was an order of magnitude higher compared to H3K4me3-negative promoters, and the signal spread over several hundred bases around nucleosomes carrying H3K4me3 (Figure S2B, Figure 1C-D left panel). Therefore, we postulated that, while the methylation state of individual adenines is inherently noisy, integrating methylation information across neighbouring adenines may robustly distinguish H3K4me3-marked molecules, from those exhibiting background mA levels (Figure 1C-D - left panel, Figure S2A-B, see Methods). To validate this approach, we quantified the single-molecule frequency of H3K4me3 marking at promoters genome-wide, and compared it to ChIP-seq enrichment (Figure S2A, see Methods). The strong correlation between these two measures suggests that our quantification strategy robustly captures H3K4me3 with singlemolecule resolution (R=0.79, Figure 2A). Similar quantification for control experiments with non-specific IgG (estimating background methylation by free-diffusing Hia5) or unmodified panH3 antibody (estimating methylation levels of Hia5 when uniformly targeted across the genome) detected low levels of mA at H3K4me3-positive loci (Figure S2D), confirming the specificity of HM detection. Together, these results demonstrate that ChromSMF quantitatively captures heterogeneity in HM deposition at single-molecule resolution.

**Fig. 2.**
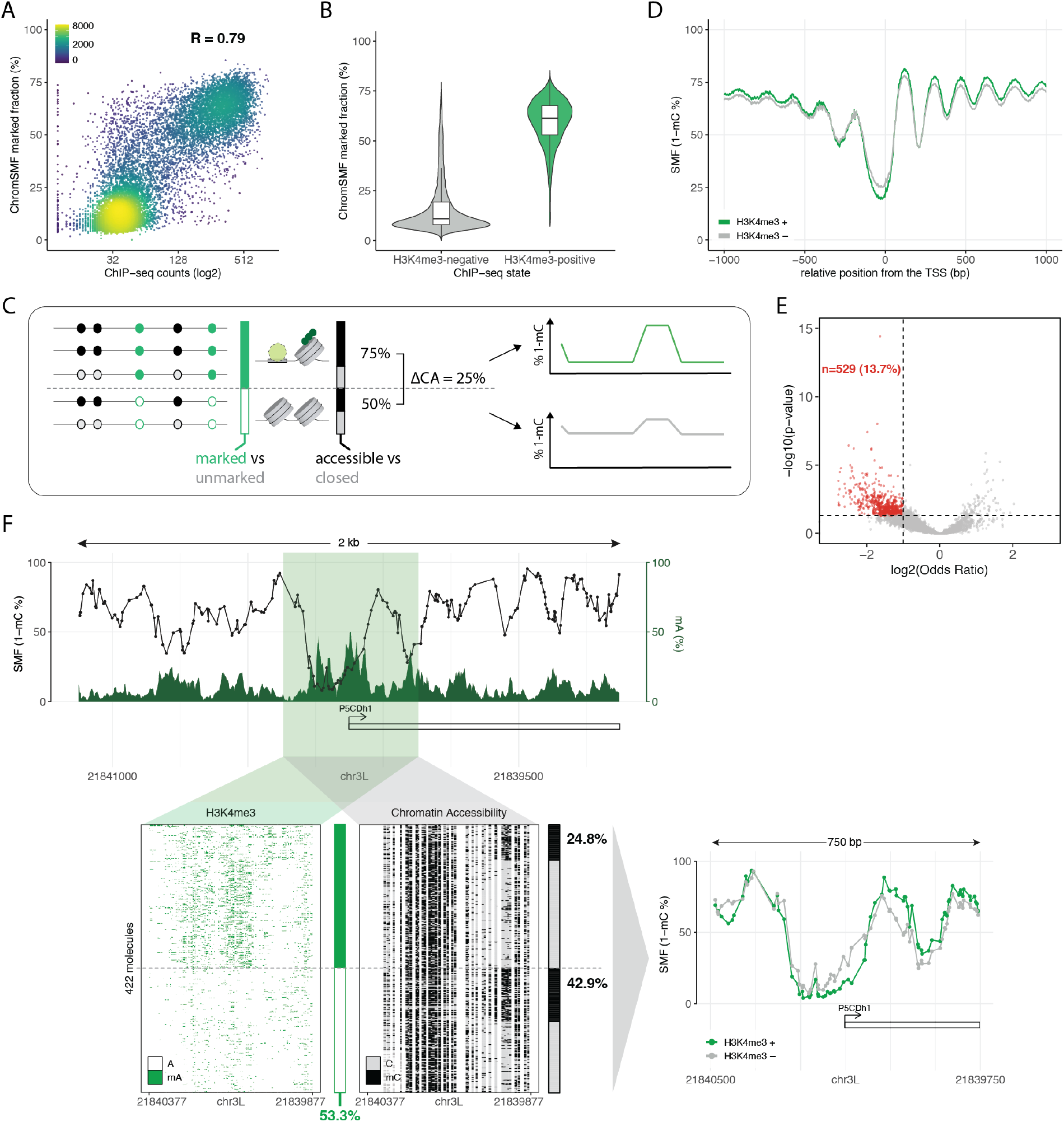
Measuring H3K4me3 and chromatin accessibility association at single-molecule resolution. **(A) Benchmarking of the single-molecule quantification of H3K4me3**. Scatter plot comparing the fraction of molecules marked by H3K4me3 as defined by ChromSMF (y-axis) against the ChIP-seq counts (log_2_) of H3K4me3 at the same genomic tile (x-axis). Data were collected from each 500 bp tile spanning at least one TSS. *R* represents the Pearson’s correlation coefficient. **(B) Distribution of the fraction of marked molecules between H3K4me3-positive and negative TSSs**. Comparison of the distribution of the fraction of marked molecules between H3K4me3-positive (green) and negative (grey) TSSs as defined by ChIP-seq. Most of the H3K4me3-positive TSSs have 50–75% H3K4me3-marked molecules as defined by the single-molecule analysis of ChromSMF data. **(C) Outline of the analysis strategy to detect histone modification–chromatin accessibility associations**. The mean mA is computed for each molecule in the extraction window, and those with mA ≥ 10% are classified as marked. For the marked and unmarked molecules, the mean chromatin accessibility (mC) is computed and molecules with mC ≥ 50% are classified as accessible. The difference in chromatin accessibility between the two fractions (marked – unmarked) is calculated and used for statistical testing of the imbalance between them using a two-sided Fisher’s exact test. **(D) H3K4me3-marked molecules have higher accessibility and stronger nucleosome positioning at TSSs**. Composite chromatin accessibility profile (1 – mC%) at H3K4me3-positive TSSs (classification based on ChIP-seq) comparing molecules classified as either marked (green) or unmarked (grey) by H3K4me3 (classification based on mA signal). **(E) Identification of TSSs with a significant association between +1 nucleosome phasing and H3K4me3**. Volcano plot depicting the odds ratio (log_2_OddsRatio; x-axis) and the p-value (− log_10_pValue; y-axis) of a two-sided Fisher’s exact test for differential chromatin accessibility between H3K4me3-marked and unmarked fractions at the same TSS. P-values were adjusted using the Benjamini–Hochberg method. Most TSSs show no chromatin accessibility–H3K4me3 association (grey dots), yet 529 TSSs show lower accessibility (higher +1 nucleosome phasing) in the H3K4me3-marked fraction (red dots). **(F) H3K4me3 is associated with higher +1 nucleosome phasing at the P5CDh1 promoter**. Top panel: ChromSMF signal for H3K4me3. Average SMF signal (1 – mC%) of individual cytosines (black) and average smoothed mA signal (mA%; green; smoothing across 4 adenines). Bottom panel - left: single-molecule stacks displaying either mA-H3K4me3 (green, left) or mC-chromatin accessibility (black, right) signal. Molecules are displayed in identical order in both panels and originate from the same sample. Single-molecule classification of H3K4me3 (green) and chromatin accessibility (black) are shown as stacked bar plots on the right of each stack. Single-molecule quantification of total H3K4me3 at the locus is shown at the bottom. Single-molecule classification of chromatin accessibility for H3K4me3-marked and H3K4me3-unmarked molecules is shown on the right. Bottom panel - right: average SMF signal (1 – mC%) of individual cytosines for molecules classified as either marked (green) or unmarked (grey) by H3K4me3.

### Measuring associations between histone modifications and chromatin accessibility at the molecular level

A unique feature of ChromSMF is that it detects HM and CA simultaneously, allowing to study their cooccurrence at the molecular level (Figure 1C-D). To that end, we next developed the computational tools to quantify coocurence of HM and CA on individual molecules. Most active promoters showed substantial epigenetic heterogeneity, with a median of 60% of molecules detected as marked by H3K4me3 (Figure 2B). We note here that absence of mA marking cannot be unambiguously interpreted as absence of H3K4me3, due to potential incomplete antibody targeting, or thetering of inactive Hia5 proteins. However, positively marked molecules can be interpreted with higher confidence. We therefore focused our analyses on these molecules, ensuring that conclusions are not confounded by false-negatives in HM labeling.

For each locus where 50-75% of the molecules were marked by H3K4me3, we computationally separated H3K4me3-marked molecules and tested if they differed in their chromatin accessibility and nucleosome occupancy profiles from unmarked ones (Figure 2C; see Methods). We observed that presence of H3K4me3 is globally associated with increased CA upstream, and increased nucleosome positioning downstream of the TSS (Figure 2D). Moreover, with a coverage of more than 500x per replicate, we had sufficient power to quantify these differences at individual promoters. We observed that H3K4me3 presence correlates with differences in chromatin organisation at about 14% of all analysed promoters (Figure 2E). These locus-specific effects can be observed when comparing individual loci. For instance, at the P5CDh1 promoter, H3K4me3-marked molecules show higher accessibility upstream and increased nucleosome phasing downstream of the TSS compared to the unmarked ones (Figure 2F). In contrast, we did not observe such H3K4me3-dependant differences in chromatin organisation at the CG9886 promoter (Figure S2E). These differences in chromatin accessibility between H3K4me3-marked and unmarked molecules were also reproducible between biological replicates (R=0.57, Figure S2F). Altogether, this demonstrates that ChromSMF captures cell-to-cell heterogeneity in HM deposition together with differences in chromatin organization, enabling direct associations between them *in vivo*.

### Molecular association of five histone modifications with chromatin accessibility

We next evaluated the applicability of ChromSMF to other HMs by generating highcoverage, genome-wide datasets for two additional active (H3K27ac, H3K4me1) and two repressive HMs (H3K27me3, H3K9me3) (median coverage 150-750x, Table S1). As expected, total mC levels, reflecting CA, were consistent across samples (Figure S3A), whereas mA distribution was specific to the targeted HM (Figure S3B). Single-molecule quantification of chromatin accessibility was likewise reproducible between samples (Figure S3C). Inspection of individual loci enriched for H3K4me1 (Figure 3A; light-green tracks), or H3K4me3 and H3K9me3 (Figure 3B; dark-green and brightred tracks) by ChIP-seq showed concordant mA marking in the corresponding ChromSMF samples, confirming specific antibody-targeting by the mA MTase. In contrast, mC-based accessibility profiles for the entire population recapitulated DNase-seq ones, and were indistinguishable across experiments performed with different antibodies (Figure 3A-B; black tracks), confirming that targeted mA deposition did not affect CA measurement. Genome-wide, single-molecule quantification of HMs also recapitulated known HM relationships: HMs associated to active enhancers, such as H3K27ac and H3K4me1, showed strong concordance (R = 0.84, Figure S3D), while the repressive H3K9me3 was largely exclusive with the active H3K4me3 (R = 0.078, Figure S3D). Furthermore, loci enriched for each HM by ChIP-seq exhibited increased single-molecule marking frequency in the corresponding ChromSMF dataset (Figure S3E-I). Together, these results demonstrate that ChromSMF can robustly profile HMs with diverse genomic distributions, while simultaneously measuring CA within the same experiment.

**Fig. 3.**
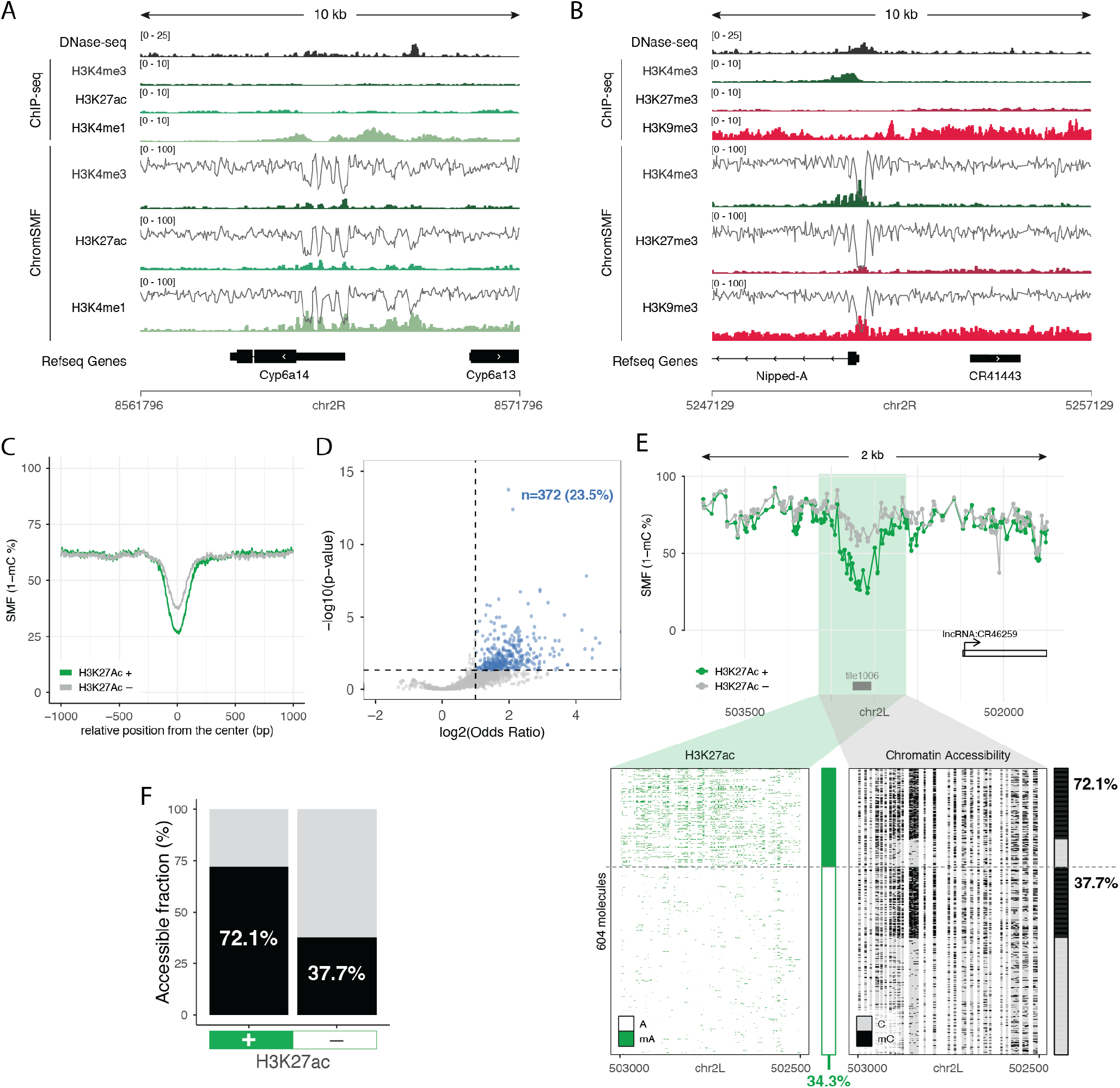
ChromSMF against five histone modifications. **(A) Measuring H3K27ac and H3K4me1 by ChromSMF**. Genome browser tracks displaying DNase-seq, ChIP-seq, and average ChromSMF signal (mA% and 1 – mC%) targeting different histone modifications associated with active regulatory elements. The genomic location between antibody-directed mA signal obtained by ChromSMF and ChIP-seq enrichment profiles shows good agreement for each modification. The chromatin accessibility profile across the entire population is comparable between experiments and in agreement with DNase-seq. **(B) Measuring H3K27me3 and H3K9me3 by ChromSMF**. Same representation as Figure 3A. **(C) H3K27ac-marked molecules show higher accessibility at enhancers**. Composite mC signal (1 – mC%) at active enhancers comparing molecules classified as either marked (green) or unmarked (grey) by H3K27ac (classification based on mA signal). **(D) Identification of enhancers with a significant association between chromatin accessibility and H3K27ac**. Volcano plot depicting the odds ratio (log_2_OddsRatio; x-axis) and the p-value (− log_10_pValue; y-axis) of a two-sided Fisher’s exact test for differential chromatin accessibility between H3K27ac-marked and unmarked fractions at the same enhancer. P-values were adjusted using the Benjamini–Hochberg method. Most enhancers show no chromatin accessibility–H3K27ac association (grey dots), yet 372 enhancers show higher accessibility in the H3K27ac-marked fraction (blue dots). **(E) H3K27ac is associated with higher chromatin accessibility at the enhancer upstream of the CR46259 gene**. ChromSMF signal for H3K27ac. Top panel: average SMF signal (1 – mC%) of individual cytosines for molecules classified as either marked (green) or unmarked (grey) by H3K27ac. Bottom panel: single-molecule stacks displaying either mA-H3K27ac (green, left) or mC-chromatin accessibility (black, right) signal. Molecules are displayed in identical order in both panels and originate from the same sample. Single-molecule classification of H3K27ac (green) and chromatin accessibility (black) are shown as stacked bar plots on the right of each stack. Single-molecule quantification of total H3K27ac at the locus is shown at the bottom. Single-molecule classification of chromatin accessibility for H3K27ac-marked and H3K27ac-unmarked molecules is shown on the right. **(F) Single-molecule quantification of chromatin accessibility at the enhancer upstream of the CR46259 gene**. Bar plot depicting the single-molecule quantification of chromatin accessibility for molecules classified as marked (left) or unmarked (right) by H3K27ac at the same enhancer.

To illustrate the power of this approach, we focused on H3K27ac, a modification that has been correlatively associated with the activation of enhancers. Consistently, we observed that, at active enhancers, DNA molecules marked by H3K27ac exhibit higher CA than the unmarked ones (Figure 3C). About 24% of the tested enhancers showed a significant association between presence of H3K27ac and increased CA at the molecular level (Figure 3D). This is for instance visible at the enhancer located upstream of the lncRNA-CR46259 (Figure 3E), where H3K27ac-marked molecules show 1.9 fold higher CA relative to unmarked ones (Figure 3E-F). In contrast, for the majority of enhancers no significant association between H3K27ac and CA was detected (Figure 3D; exemplified in Figure S3J-K). Altogether, this illustrates the ability of ChromSMF to robustly capture context-specific associations between various HMs and CA at diverse *cis*-regulatory elements.

### Identification of long-range associations between histone modifications and chromatin accessibility

The median read length in our ChromSMF datasets is 5 kilobases (Table S1). Therefore, individual molecules often span multiple cREs while providing continuous information on CA and HM (Figure S4A). We therefore extended our computational strategy to measure molecular associations over longer distances, linking the presence of a HM at a given cRE, with CA at other cREs present in *cis* (Figure 4A, see Methods). Using this approach, we tested whether the presence of H3K27ac at enhancers is associated with increased chromatin accessibility at surrounding enhancers and promoters. We identified a subset of enhancers for which the positive molecular association between H3K27ac and CA was not restricted to the marked enhancer itself but extended to neighboring cREs (Figure 4B). Moreover, these associations were independent on the distance from the viewpoint enhancer (Figure 4B), and were observed at both distal promoters and enhancers (Figure S4B). In contrast, we did not observe distal effects for H3K4me3. Instead, H3K4me3 was associated with differences in CA and nucleosome occupancy only at the promoter where the modification was deposited, suggesting these longrange associations are specific to a subset of HMs (Figure 4C, Figure S4C-F). An example of such long-range association can be observed at the COX7AL2 gene locus, where the presence of H3K27ac at an enhancer upstream of the gene, correlates with increased accessibility at multiple neighbouring cREs, including the promoter element (Figure 4D-H). Together, these results demonstrate that ChromSMF can identify molecular associations between multiple regulatory layers across distances of up to several kilobases.

**Fig. 4.**
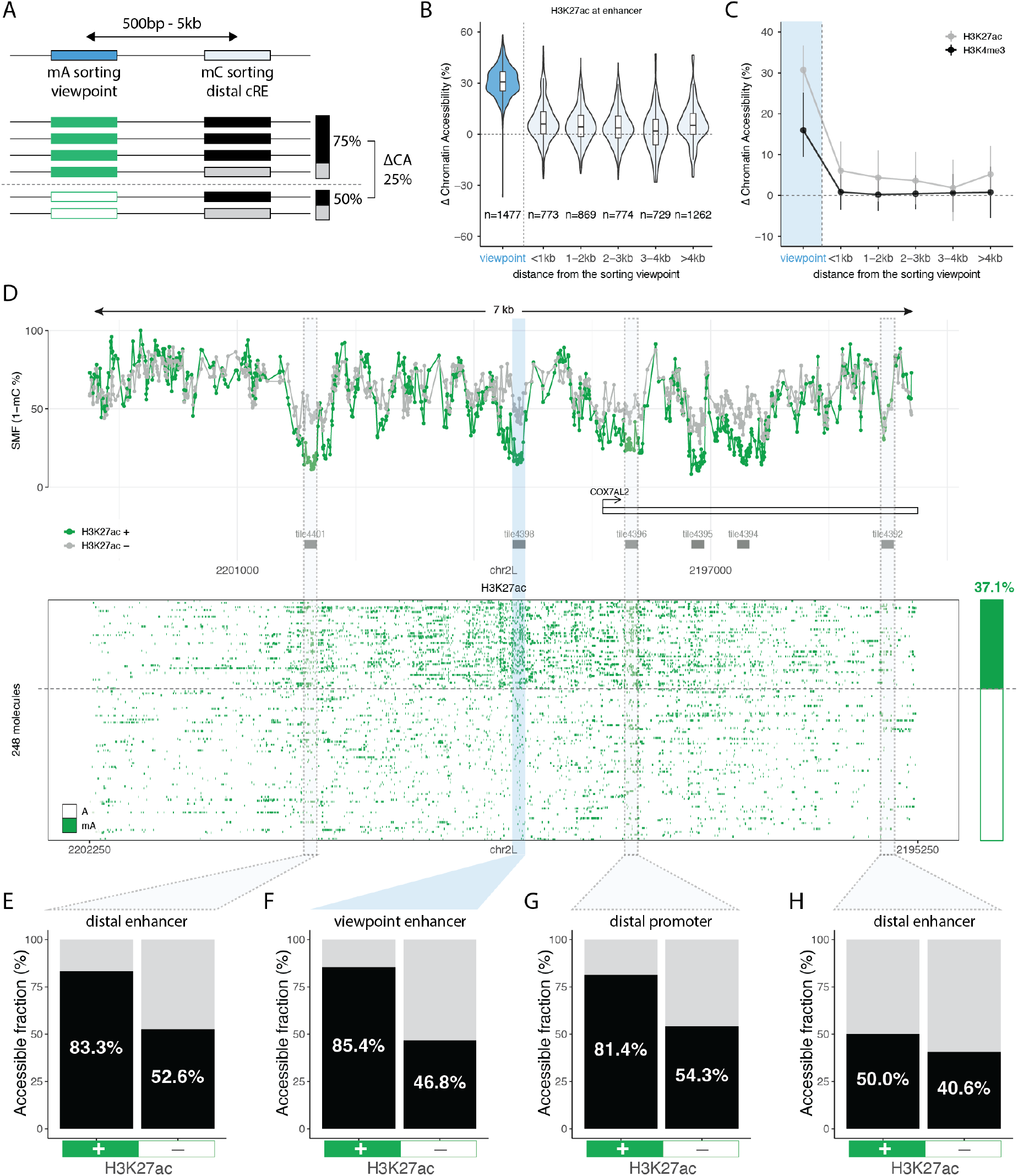
Molecular association between histone modifications and chromatin accessibility over long genomic distances. **(A) Outline of the analysis strategy to detect HM–CA associations over long genomic distances**. DNA molecules are sorted into HM-marked or unmarked based on a 500 bp window centred at a cRE (viewpoint). The chromatin accessibility differences between the two fractions (marked – unmarked at the viewpoint) is computed for all the cREs located within 5 kilobases from the viewpoint (distal cRE). The difference in chromatin accessibility between the two fractions is computed to link the presence of the HM with its potential consequences on chromatin accessibility at distant cREs. **(B) H3K27ac marking associates with chromatin accessibility differences at distant cREs**. Distribution of the difference in chromatin accessibility between H3K27ac-marked and unmarked fractions at the viewpoint (enhancers; blue) and at cREs located at various distances from it (white). **(C) H3K4me3 marking is not associated with chromatin accessibility differences at distant cREs**. Comparison of the long-range molecular association between histone modifications and chromatin accessibility for H3K27ac (grey) and H3K4me3 (black). Dots represent the median value, and vertical lines represent the interquartile range (q25–q75) for each distribution. The blue shaded area highlights the CA delta for the viewpoint. **(D) Single locus example where H3K27ac at an enhancer is associated with higher chromatin accessibility at multiple cREs of the *cis*-regulatory landscape**. Top panel: ChromSMF signal for H3K27ac. Average SMF signal (1 – mC%) of individual cytosines from molecules classified as either marked (green) or unmarked (grey) by H3K27ac. Bottom panel: single-molecule stacks displaying mA-H3K27ac (green) signal. Single-molecule classification of H3K27ac at the viewpoint enhancer is shown as a stacked bar plot on the right. Single-molecule quantification of total H3K27ac abundance at the viewpoint enhancer (green) is shown on top of the bar plot. **(E–H) H3K27ac associates with increased chromatin accessibility at a subset of distal cREs**. Bar plot depicting the single-molecule quantification of chromatin accessibility for molecules classified as H3K27ac-marked (left) or unmarked (right) at the enhancer upstream of the COX7AL2 promoter. CA is quantified at the enhancer itself **(F)**, at the nearby COX7AL2 promoter **(G)**, or at other enhancers in the locus **(E, H)**.

### Allele-specific epigenetic regulation in mammalian genomes

Having established a robust protocol in *Drosophila*, we next tested the portability of ChromSMF to study gene expression regulation in mammals. This presents additional challenges due to larger genomes, a more complex epigenetic landscape which includes endogenous DNA methylation, and genetic variation between individuals and alleles. To benchmark the ability of ChromSMF to capture the complex crosstalk between genetic and epigenetic regulation in an allele-specific manner, we used a mouse embryonic stem cell (mESC) line derived from the F1 cross between two evolutionary distant mouse species (*Mus musculus domesticus* and *Mus musculus castaneus* (Bl6xCast)) (51) (Figure 5A). We performed ChromSMF against H3K4me3, and sequenced it to 30x (Table S2), a coverage that can be obtained using a single PromethION flow cell (typical output above 100 Gb). This coverage was sufficient to simultaneously quantify H3K4me3, CA, and 5mC. Moreover, as genotype information is contained within individual sequencing reads, signals could be assigned to each allele and directly compared (Figure 5A, Figure S5A). To illustrate the power of this multi-channel allele-specific analysis, we focused on imprinted control regions (ICRs), where 5mC establishes parentally inherited monoallelic gene expression, and where 5mC and CA are expected to occur on opposite alleles (52). Consistently, ChromSMF recapitulated known allelic differences in 5mC at ICRs (Figure S5B). As an example, we looked at the *Kcnq1ot1* locus, a paternally expressed ICR (Figure 5B-F). In addition to differential 5mC (Figure 5C, F), we observed differences in CA (Figure 5D, F), and H3K4me3 (Figure 5B, F) that are primarily found on the unmethylated paternal promoter, and absent from the repressed maternal allele (Figure 5B-F). In contrast, non-ICR control promoters showed comparable levels of 5mC, H3K4me3 and CA on both alleles (Figure S5C-E). Inspection of allele-specific CA at the *Kcnq1ot1* locus, we noticed a short region within the gene body that showed differential accessibility between alleles, a pattern typical of active enhancers (23). This was at first surprising as the higher accessibility occurred on the not accessible, repressed, maternal allele (Figure 5D). More detailed inspection of the CA profile revealed two short footprints that we hypothesised to be created by the binding of TFs (Figure 5G), as this can be measured and quantified by SMF at single-molecule resolution (24, 53). Screening for TF binding motifs, we found characteristic motif for the ZFp57 (Zinc finger protein 57), a methylation-dependant zinc-finger TF involved in heterochromatin-driven gene repression (Figure 5G) (54). Cross-validation with ChIP-seq data confirmed ZFp57 binding at this repressor element (Figure 5G). Together, this shows that ChromSMF can simultaneously quantify HM, CA, endogenous 5mC and TF binding, at allelic resolution. Moreover, it illustrates how this information can be combined to resolve complex allele-specific epigenetic regulation in mammalian genomes.

**Fig. 5.**
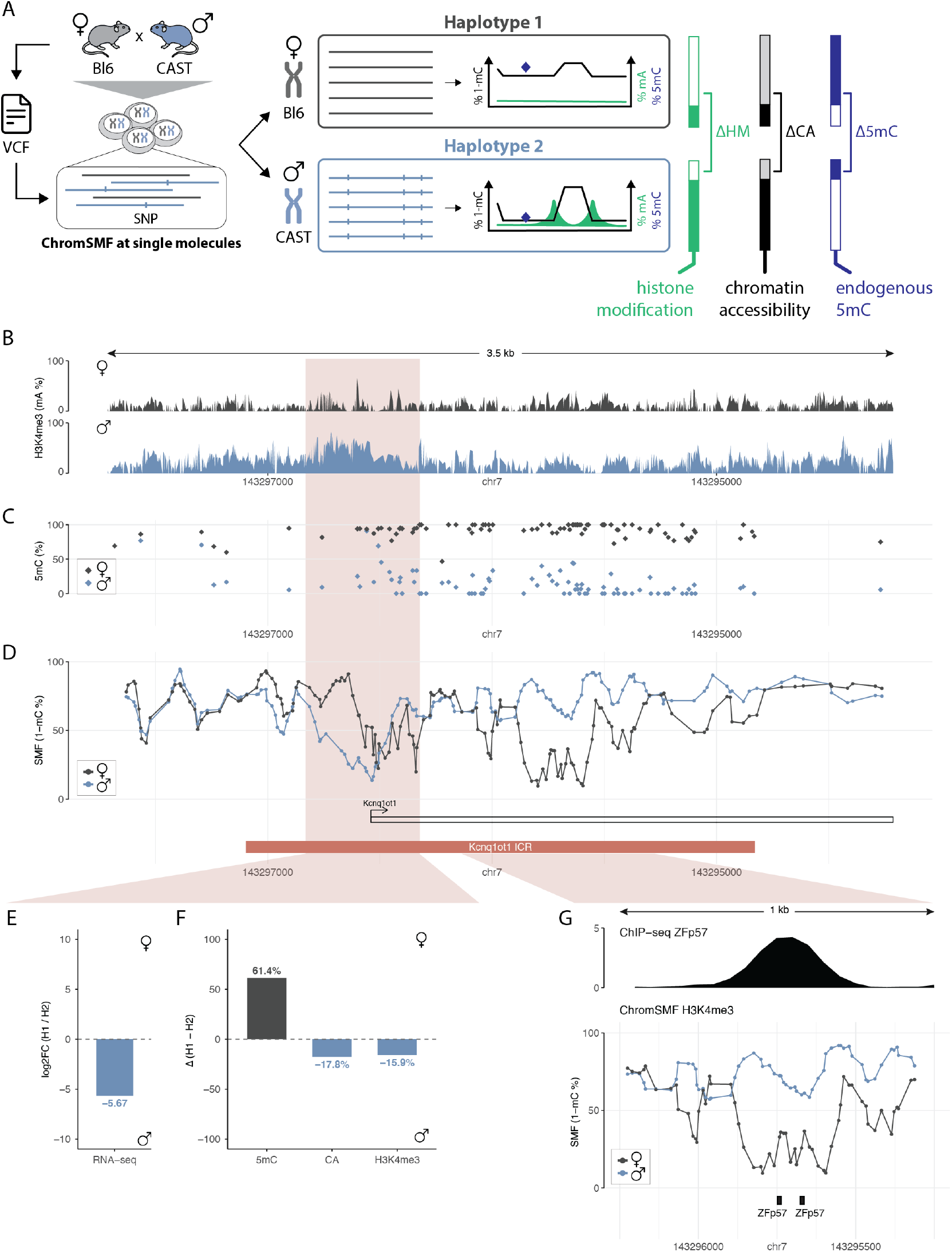
Resolving multiple epigenetic layers on individual alleles. **(A) Outline of the analysis strategy to simultaneously measure histone modifications, chromatin accessibility, and DNA methylation on individual alleles**. Long reads are mapped to individual alleles using reference Single Nucleotide Polimorfisms (SNPs) between strains (Bl6/CAST). Differences between alleles are then computed for each channel: 5mC (blue diamond), chromatin accessibility (CA; black SMF track), and H3K4me3 levels (HM; green track). **(B–D) Parallel analysis of multiple epigenetic layers on individual alleles at the *Kcnq1ot1* imprinted control region**. ChromSMF was performed in F1 hybrid (Bl6/CAST) mESCs targeting H3K4me3. Signals are plotted for the two alleles separately and the colour code reflects haplotype origin: Haplotype1 (H1, ♀, maternal) is represented in dark-grey, and Haplotype2 (H2, ♂, paternal) is represented in blue. All signals originate from the same sample. **(B)** Average mA signal (mA%) smoothed every 3 adenines within a 3.5 kb window around the annotated ICR. **(C)** Average 5mC signal (mC%) at individual CpGs for the same region. **(D)** Average SMF signal (1 – mC%) at the same region, smoothed every 4 GpCs to compensate for low coverage. The ICR position is highlighted with a brick-red rectangle. **(E–F) Quantification of allelic differences at the *Kcnq1ot1* promoter**. Bar plots showing differences between alleles (computed as H1 – H2) at the *Kcnq1ot1* promoter for **(E)** gene expression (log_2_ FoldChange; RNA-seq), as well as **(F)** endogenous methylation (5mC; average mC% in CpG context), chromatin accessibility (CA; average mC% in GpC context), and H3K4me3 (single-molecule sorting for H3K4me3-mA using a 100 bp window around the TSS). Colours depict which allele shows the higher value (maternal = dark-grey, paternal = blue). Endogenous 5mC is higher in the repressed allele ( ♀, maternal), whereas chromatin accessibility and H3K4me3 are higher in the expressed allele ( ♂, paternal). **(G) ChromSMF detects footprints for the repressor Zfp57 on the repressed allele**. Top panel: ChIP-seq track for Zfp57. Bottom panel: representation as in (D) zoomed in to 1 kb window in the gene body. Zfp57 motifs are highlighted as black rectangles.

### Haplotype-resolved analysis of epigenetic mechanisms contributing to allelic imbalance

Because ChromSMF uses long-read sequencing, it is compatible with allele-specific epigenomic analyses in samples lacking a reference genome, a common situation in clinical settings. We therefore adapted our computational framework to enable data-driven haplotype reconstruction without prior knowledge of allele-specific phase blocks (55, 56). To benchmark this, we performed haplotype separation of the H3K4me3 ChromSMF data from F1 mESC (Bl6/CAST), for which the genetic ground truth is known (57). Across autosomes, we resolved 95% of the genome into 7123 phase blocks with a median length of 28 kb, while only 18% of the reads remained unassigned (Figure S6A-C). To illustrate the power of the approach, we examined whether we could identify epigenetic mechanisms associated with allele-specific gene expression (Figure S6D). Excluding ICRs, differential DNA methylation was detected at only 12% of the imbalanced genes (Figure 6B). A slightly higher proportion of promoters showed differences in CA (Figure 6B), and about one quarter exhibited allele-specific enrichment of H3K4me3 (Figure 6B). Taken either individually or in combination, these three layers accounted for 42% of the observed allelic imbalance in gene expression (Figure S6E). The specificity of this allele-resolved epigenetic detection is exemplified at the *Tal2* promoter (Figure 6C-E), where the frequency of H3K4me3 marking is 36% higher on the allele with higher expression (Log2(FC) = 4.28; Figure 6D-E). In contrast, both alleles show comparable levels of CA and 5mC (Figure 6E). Such disparities would have been missed by existing approaches that measure CA or 5mC in isolation (26), underscoring the importance of integrating HM information to explain allelic differences in gene expression. More broadly, this demonstrates the potential of ChromSMF for allele-resolved dissection of epigenetic regulation of gene expression in complex genomes, extending beyond model organisms to include human and clinical samples.

**Fig. 6.**
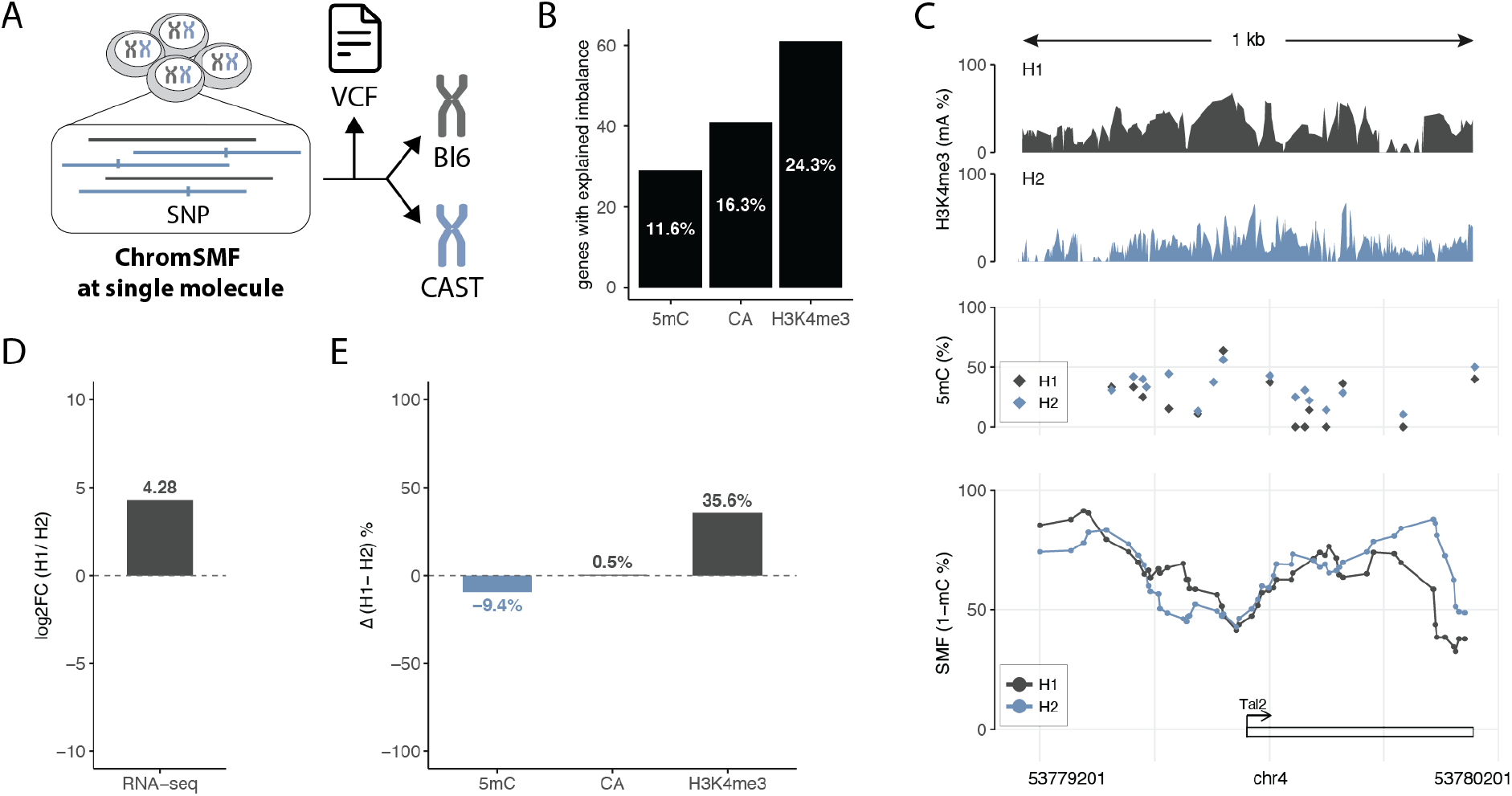
Simultaneous haplotype-resolved analysis of multiple epigenetic layers. **(A) Outline of the analysis strategy for reference-free haplotype-resolved analysis of multiple epigenetic layers**. Long reads are directly phased into haplotypes without prior knowledge of genetic variation between strains, and haplotype-specific signal is resolved for H3K4me3, chromatin accessibility, and DNA methylation. **(B) Systematic comparison of the association of each epigenetic layer with allelic imbalance in gene expression**. Bar plot showing the number and proportion of genes with significant allelic imbalance in RNA-seq (| log_2_ FoldChange| ≥ 1 and adjusted p-value *<* 0.05) that also exhibit haplotype asymmetry in DNA methylation (5mC; Δ ≤ −0.15), chromatin accessibility (CA; Δ ≥ 0.15), or H3K4me3 signal (HM; Δ ≥ 0.15). Deltas are computed as (highlyExpressed allele – lowlyExpressed allele). Percentages (in-bar labels) are calculated relative to the total number of genes with allelic imbalanced expression that were analysed (*n* = 251). **(C–E) Allelic differences in H3K4me3 but not in chromatin accessibility associate with allelic imbalance of the *Tal2* gene. (C)** Average ChromSMF tracks within a 1 kb window around the TSS. All signals originate from the same sample. Colour code reflects haplotype assignment (haplotype 1 = dark-grey, haplotype 2 = blue). Top panel: average mA signal (mA%) smoothed every 3 adenines. Middle panel: average 5mC signal (mC%) at individual CpGs. Bottom panel: average SMF signal (1 – mC%) smoothed every 4 GpCs to compensate for low coverage. **(D)** Bar plot showing the fold change (log_2_ FC) of *Tal2* expression between haplotypes. **(E)** Bar plot showing haplotype asymmetry at the *Tal2* promoter for endogenous methylation (5mC; average mC% in CpG context), chromatin accessibility (CA; average mC% in GpC context), and H3K4me3 (single-molecule sorting for H3K4me3-mA using a 100 bp window around the TSS). Colours depict which allele shows the higher value (haplotype 1 = dark-grey, haplotype 2 = blue). While 5mC and CA show small differences between alleles, the single-molecule H3K4me3 quantification shows a delta of 35.6%, indicating a higher fraction of H3K4me3-marked molecules in the highly expressed allele (haplotype 1).

## Discussion

Here, we developed and benchmarked ChromSMF, a long-read single-molecule genomic method for the integrated analysis of up to five layers of genetic and epigenetic information. Unlike enrichment-based assays, ChromSMF preserves cell-to-cell heterogeneity and enables direct quantification of the frequency of chromatin states. By leveraging continuous, multi-kilobase information, ChromSMF allows the study of the combinatorial occurrence of histone modifications, chromatin accessibility, TF binding, and DNA methylation on the same DNA molecules. This allows direct interrogation of relationships between multiple regulatory layers to identify mutually exclusive, cooperative, or locus-specific associations that are otherwise inaccessible to bulk assays. We anticipate ChromSMF will be transformative for time-course studies of chromatin regulation, enabling dissection of the temporal order of TF binding and histone modification changes during cell-state transitions in development or disease.

We demonstrated the robustness and portability of ChromSMF across multiple histone modifications and model systems. Beyond histone modifications, the approach is, in principle, applicable to a wide range of chromatin-associated proteins for which antibodies exist. For instance, we anticipate ChromSMF to be applicable to TFs, histone variants, RNA polymerase II and its phosphorylation states, and other regulators of genome function (40–44). Moreover, the use of distinct DNA-modifying enzymes pre-incubated with different antibodies could increase the number of genomic features measurable on the same DNA molecule, further expanding the multiplexing capacity of the method.

The continuity of ChromSMF data over multiple kilobases enables the study of coordinated regulatory states across promoters, gene bodies, and distal *cis*-regulatory elements. Extended read lengths further facilitate mapping in repetitive or structurally complex genomic regions, expanding the range of regulatory contexts and genomes that can be interrogated. By jointly measuring sequence variation and multiple epigenetic layers on individual molecules, ChromSMF directly links genotype, chromatin state, and regulatory outcome. More broadly, it provides a mechanistic framework to connect disease-associated mutations with epigenomic alterations and their functional consequences on transcriptional regulation. Moreover, ChromSMF data can be directly phased to resolve haplotypes, making it applicable in clinical settings in which personalized genome assemblies are not available.

In its current design, ChromSMF requires approximately 1 µg of DNA (10^5^ mammalian cells), considerably more than low-input enrichment-based assays (*i*.*e*., CUT&RUN). While this could potentially be reduced, it would likely compromise coverage, an essential parameter for the robustness of the assay. The experimental protocol can be completed within approximately one week. Sample preparation is relatively inexpensive (around 200 EUR) and can be performed with conventional laboratory equipment by researchers with standard molecular biology training. However, because ChromSMF does not rely on enrichment, sequencing requirements are higher than for conventional bulk assays (typically one PromethION flow cell per mammalian sample). While PCR-based enrichments are not compatible with the method, future adaptation of PCR-free enrichment strategies for long-read technologies (*e*.*g*., Nanopore adaptive sampling (58, 59)) should mitigate sequencing costs for applications that do not require genome-wide coverage. Finally, although primarily implemented using ONT, ChromSMF is expected to be also compatible with PacBio HiFi sequencing. This may be advantageous for applications requiring highly accurate base calling, albeit at higher per-base cost.

Together, ChromSMF provides a unified and quantitative framework for dissecting combinatorial genetic and epigenetic states that underlie transcriptional regulation in healthy and disease contexts.

## Methods

### M.CviPI protein purification

Purification of the M.CviPI protein (pBAD-HisMBP3C-McviPI) was performed as previously described (23). For the detailed and updated purification protocol, refer to the following: https://dx.doi.org/10.17504/protocols.io.eq2ly776plx9/v1.

### pA-Hia5 protein purification

Purification of the pA-Hia5 fusion protein (pET-pA-Hia5; Addgene, 174372) was performed as previously described (40). For the detailed and updated purification protocol, refer to the following: https://dx.doi.org/10.17504/protocols.io.bv82n9ye.

### pA-Hia5 activity assay *in vitro*

Hia5 activity of the purified pA-Hia5 protein was assayed *in vitro* as follows. Naked unmethylated Lambda DNA (Promega, D152A) was methylated by a range of pA-Hia5 concentrations (500nM, 250nM, 125nM, 62.5nM, 31.25nM, 15.63nM, 0nM). The DNA was subsequently digested by DpnI, which can cut Gm6ATC but not GATC sequences and analyzed on agarose gel. In summary, serial dilutions of pA-Hia5 were made using dilution buffer A (25 mM Tris pH 8.0, 50 mM NaCl, 50% glycerol). For each concentration, 1 µg of DNA was incubated with 14 µl of Activation Buffer (15 mM Tris, pH 8.0, 15 mM sodium chloride, 60 mM potassium chloride, 1 mM EDTA, pH 8.0, 0.5 mM EGTA, pH 8.0, 0.5 mM spermidine (Sigma, S0266-5G), 0.1% BSA (Sigma, A2153-10G)), 3.2 mM SAM (2 µl, NEB, B9003S), and 2 µl of the desired dilution of pA-Hia5 protein for 1 hour at 37°C. Next the reaction mix was heat inactivated for 15 min at 65°C, and 15 µl of it was mixed with 1x rCutSmart buffer (NEB, B6004S) supplemented with 10 units of DpnI enzyme (NEB, R0176S) for 1 hour at 37°C and run on an agarose gel to assess the degree of methylation. As a comparison and positive control, we performed the same assay in parallel using an aliquot of pA-Hia5 from the Altemose Lab. We confirmed methylation activity of the purified pA-Hia5 was comparable to the reference one, with a saturation point around 25 nM. We therefore used the same pre-tested amount (200nM, saturating *in vitro*) for all our experiments.

### Cell culture *Drosophila* cell line

*Drosophila* embryonic S2 cells were cultured at 25°C (without CO2) on 15cm plates in Schneider’s *Drosophila* medium (Gibco, 21720024) supplemented with 10% of inactivated Fetal Bovine Serum (FBS). Cells were split every 3 to 5 days at 2 million cells per mL for a maximum of 10 passages.

### Cell culture mouse cell line

Mouse F1 hybrid ES cells (male Bl6/CAST (51)) were cultured at 37°C and 5% CO2 on 0.2% gelatin-coated 10cm plates in ES medium (Dulbecco’s Modified Eagle Medium (DMEM), supplemented with 15% heat inactivated Fetal Bovine Serum (FBS), 20ng/mL Leukemia Inhibitory Factor (LIF), 0.05 mM 2-Mercaptoethanol, 1 mM Sodium pyruvate, 2 mM LGlutamine and 1x non-essential amino acids (NEAA)). Cells were trypsinized and split every second day for a maximum of 10-15 passages.

### Immobilization on ConA beads and Digitonin permeabilization

All reagents were prepared fresh, and kept on ice. To equilibrate beads, Concanavalin A bead slurry (ConA, Polysciences Europe GmbH, 86057-3) were resuspended by gentle vortexing and 20 µL per condition was added to 1.5 mL ConA binding buffer (20 mM HEPES pH 7.5, 10 mM KCl, 1 mM CaCl2, 1 mM MnCl2; stable at 4°C for 6 months), gently resuspended and placed on a magnet. Supernatant was discarded, and the washing step was repeated again. Washed beads were resuspended in 20 µL ConA binding buffer per condition. Cells were pelleted at 500-1000g for 5 min at 4°C and washed with PBS twice. Pelleted cells were resuspended in 1 mL Wash buffer per condition (20 mM HEPES-potassium hydroxide buffer pH 7.5, 150 mM sodium chloride, 0.5 mM Spermidine (Sigma, S0266-5G), and 1 Roche cOmplete EDTA-free tablet (Merck, 11873580001) per 50 mL buffer) and split into separate eppendorf’s tubes. 20 µL of equilibrated ConA beads were added to each tube, incubated at RT for 10 min (optional 500 rpm), then placed on a magnet. Supernatant was removed, cells were resuspended in 1 mL ice-cold Dig-Wash buffer (0.04% digitonin (Sigma, 300410-250MG), 20 mM HEPES-potassium hydroxide buffer pH 7.5, 150 mM sodium chloride, 0.5 mM Spermidine (Sigma, S0266-5G), 1 Roche cOmplete EDTA-free tablet (Merck, 11873580001) per 50 mL buffer, and 0.1% BSA) and incubated on ice for 5 min. Digitonin concentration must be adjusted depending on the cell type and the stock solution. The suspension was then placed on a magnet, supernatant was discarded, and cells were resuspended in the specific condition solution. For SMF and ChromSMF protocols, cells were resuspended in 94.5 µl of 1X M.GpC buffer (NEB, B0227S). For DiMeLo-seq protocol cells were resuspended in 200 µl of Tween-wash buffer (0.1% Tween-20 (Sigma, P1379), 20 mM HEPES-potassium hydroxide buffer pH 7.5, 150 mM sodium chloride, 0.5 mM Spermidine (Sigma, S0266), 1 Roche cOmplete EDTA-free tablet (Merck, 11873580001) per 50 mL buffer, and 0.1% BSA (Sigma, A2153)) supplemented with the specific antibody. Antibody concentration depends on the antibody and target of interest (Table S3).

Number of cells per condition was enough to get 1 µg of genomic DNA (*e*.*g*., 3*x*10^6^ cells for *Drosophila*, 0.25*x*10^6^ cells for mouse). Cell viability at harvesting was > 80% for *Drosophila* and > 95% for mouse experiments. Permeabilization was assessed after 5 min incubation with digitonin using trypan blue staining, and for all experiments this was >80% (<20% live cells).

### ChromSMF: Chromatin-informed Single Molecule Footprinting

All reagents were prepared fresh, and kept on ice. Cells were collected, pelleted, washed twice with PBS, immobilised on ConA beads and permeabilised with Digitonin. Permeabilised cells were resuspended in 94.5 µl 1X M.GpC buffer (NEB B0227S). For the GpC methylation treatment, 150 µl GpC methyltransferase mix (1x M.GpC buffer, 450 mM sucrose, 0.96 mM SAM (NEB, B9003S)) and 200 U M.CviPI were added and incubated at 30°C for 7.5 min. Reaction was replenished with 100 U of M.CviPI and 0.38 mM of SAM and incubated again at 30°C for 7.5 min. For the CpG methylation treatment, 0.38 mM of SAM and 40U of M.SssI (NEB, M0226L) were added to the suspension and incubated at 30°C for 7.5 min. Cells were then placed on a magnet, supernatant was removed, and washed twice with 0.95 mL of Tween-Wash (0.1% Tween-20 (Sigma, P1379), 20 mM HEPES-potassium hydroxide buffer pH 7.5, 150 mM sodium chloride, 0.5 mM Spermidine (Sigma, S0266), 1 Roche cOmplete EDTA-free tablet (Merck, 11873580001) per 50 mL buffer and 0.1% BSA (Sigma, A2153)). For each wash, the pellet was completely resuspended by pipetting up and down around ten times using wide boar tips and placed on a rotator at 4°C for 5 min before clearing on a magnet. After placing on a magnet, supernatant was discarded, and the pellet was gently resuspended in 200 µl Tween-Wash containing the primary antibody at a the desired dilution (ensuring primary antibody species was compatible with pA). Samples were placed on a rotator at 4°C overnight to allow antibody to bind. The next day, cells were placed on a magnet, supernatant was discarded, and washed twice with 0.95 mL Tween-Wash. For each wash, the pellet was completely resuspended by pipetting up and down around ten times using wide boar tips and placed on a rotator at 4°C for 5 min before clearing on a magnet. After placing on a magnet, supernatant was discarded, and the pellet was gently resuspended in 200 µl Tween-Wash containing 200 nM pA-Hia5 (ensuring pA-Hia5 activity was comparable between batches). Samples were placed on a rotator at 4°C for 2 hours to allow pA-Hia5 binding to the antibodies. Cells were then placed on a magnet, supernatant was discarded, and washed twice with 0.95 mL Tween-Wash with a 4°C rotating incubation for 5 min in between, as for the wash following antibody binding. After placing on a magnet, supernatant was discarded, and the pellet was gently resuspended in 100 µl of Activation Buffer (15 mM Tris, pH 8.0, 15 mM sodium chloride, 60 mM potassium chloride, 1 mM EDTA, pH 8.0, 0.5 mM EGTA, pH 8.0, 0.05 mM Spermidine (Sigma, S0266), 0.1% BSA (Sigma, A2153) and 800 µM SAM (NEB, B9003S)) and incubated at 30°C for 2 hours. Cells were pipet mixed every 30 min. SAM was replenished after 1 hour by adding an additional 800 µM (final concentration). Finally, the methylation reaction was stopped by adding 20 µl proteinase K (20 mg/mL), 1 µl SDS 20% and 11 µl NaCl 5 M, and incubated overnight at 55°C. The next day, DNA was extracted using the Quick-DNA HMW MagBead Kit.

Number of cells per conditions was enough to get 1 µg of genomic DNA (e.g. 3*x*10^6^ cells for *Drosophila*, 0.25*x*10^6^ cells for mouse). For mESC cells with endogenous CpG methylation, the footprinting step using M.SssI (CpG methyltransferase) was omitted, washing and antibody binding were performed directly after the second 7.5 min incubation with M.CviPI. Activation step for mESC was done at 37°C, for S2 cells at 30°C. Antibody references and concentration used are listed in Table S3.

For experiments targeting H3K27ac, 5 mM Sodium butyrate was added to PBS and all the ChromSMF buffers except for the Activation Buffer (Wash Buffer, Dig-Wash Buffer, Tween-Wash Buffer).

### DiMeLo-seq: Directed Methylation with Long-read Sequenching

DiMeLo-seq against H3K4me3 was performed as previously described with few adaptations (40, 42). In summary, all reagents were prepared fresh, and kept on ice. Around 2.5*x*10^6^ S2 cells were pelleted, washed with PBS twice, immobilised on ConA beads and permeabilised with Digitonin. Permeabilised cells were resuspended in 200 µl Tween-Wash containing the primary antibody at the desired dilution (ensuring primary antibody species is compatible with pA). Samples were placed on a rotator at 4°C overnight to allow antibody to bind. Cells were then placed on a magnet, supernatant was discarded, and washed twice with 0.95 mL Tween-Wash. For each wash, the pellet was completely resuspended by pipetting up and down around ten times using wide boar tips and placed on a rotator at 4°C for 5 min before clearing on a magnet. After placing on a magnet, supernatant was discarded, and the pellet was gently resuspended in 200 µl Tween-Wash containing 200 nM pA-Hia5. Samples were placed on a rotator at 4°C for 2 hours to allow pA-Hia5 binding to the antibodies. Cells were then placed on a magnet, supernatant was discarded, and washed twice with 0.95 mL Tween-Wash with a 4°C rotating incubation for 5 min in between, as for the wash following antibody binding. After placing on a magnet, supernatant was discarded, and the pellet was gently resuspended in 100 µl of Activation Buffer (15 mM Tris, pH 8.0, 15 mM sodium chloride, 60 mM potassium chloride, 1 mM EDTA, pH 8.0, 0.5 mM EGTA, pH 8.0, 0.5 mM spermidine (Sigma,S0266), 0.1% BSA (Sigma, A2153) and 800 µM SAM (NEB, B9003S)) and incubated at 30°C for 2 hours. Cells were pipet mixed every 30 min. SAM was replenished after 1 hour by adding an additional 800 µM (final concentration). Finally, the methylation reaction was stopped by adding 20 µl proteinase K (20 mg/mL), 1 µl SDS 20% and 11 µl NaCl 5 M, and incubated overnight at 55°C. The next day, DNA was extracted using the Quick-DNA HMW MagBead Kit.

### High Molecular Weight DNA extraction

DNA was extracted using the Quick-DNA HMW MagBead Kit (Zymo, D6060) following the manufacturer’s instructions with few adaptations. In summary, pellets were lysed in 200 µL Biofluid&SolidTissue Buffer supplemented with 20 µL Proteinase K (20 mg/mL) and 1 µL RNase A (10 mg/mL), mixed by gentle inversion, and incubated at 37°C for 1 hour at 1300 rpm in a thermomixer. When ConA beads had been used upstream, samples were placed on a magnetic rack and the supernatant transferred to a fresh low-bind tube prior to purification. An equal volume (400 µL) of Quick-DNA Mag-Binding Buffer was added, followed by 33 µL MagBinding Beads, mixed by inversion, and incubated for 10 min at room temperature at 1300 rpm. Samples were cleared on a magnet, the supernatant discarded, and beads resuspended in 500 µL Quick-DNA MagBinding Buffer, incubated for 5 min at room temperature at 1300 rpm, and cleared again. Beads were washed once with 500 µL DNA Pre-Wash Buffer and twice with 900 µL g-DNA Wash Buffer, fully resuspended during each wash and transferred to new low-bind tubes before magnetic separation. Supernatant was discarded and beads were dried at 55°C for 10 min, and DNA was eluted in 50 µL prewarmed (55°C) DNA Elution Buffer, following incubation at 55°C for at least 10 min at 1300 rpm. Wide-bore tips were used throughout mixing steps to minimize DNA shearing. Eluted DNA was used immediately for library preparation or stored at 4°C (short term).

### DNA fragmentation with gTubes

DNA fragment sizes were homogenised to 8-10 kb using g-Tubes (Covaris, 520079) following manufacturer’s instruction. In summary, samples were loaded to g-Tubes and topped up to 150 µl with water. Tubes were centrifugated twice at 5000 rpm 60 sec, twisted, and centrifugated twice at 5000 rpm 60 sec again. Full volume was retrieved in a low binding eppendorf’s tube. For mouse samples, centrifugation speed was reduced to 4200 rpm to get longer fragments (around 10-12 kb).

### DNA size quality check with FEMTO

DNA size was checked by capillar electrophoresis using the Genomic DNA 165 kb kit from Femto Pulse System (Agilent, FP-1002-0275) following manufacturer’s instruction.

### DNA quantification with 1x high sensitivity Qubit

DNA concentration was determined using the Qubit dsDNA High Sensitivity Assay Kit (Thermo Fisher Scientific) following manufacturer’s instruction.

### Nanopore Library Preparation and sequencing

Library preparation was performed using the ligation sequencing kit SQK-LSK114 following manufacturer’s instruction. 20-40 fmol of DNA were loaded in the R10.4.1 flow cell. Multiple rounds of washing and loading were performed when possible (Flow Cell Wash Kit XL, Nanopore, EXP-WSH004-XL). For each sample, we performed up to a total of 4 loads depending on quantity of input library material.

### Generation of training data for methylation calling

Training data for the methylation calling model were generated as previously described (23). In summary, naked genomic DNA (gDNA) was extracted from *Drosophila* S2 cells using the Quick-DNA HMW MagBead Kit (Zymo, D6060) and fragmented to 6 kb using g-Tubes (Covaris, 520079) following the manufacturer’s instructions. gDNA was then *in vitro* methylated using mA and mC MTases individually (sample 3 and sample 4) or in combination (sample 7). Fully methylated CpG and GpC gDNA (sample 4) was generated by two consecutive 30 min incubations at 37°C with 8 U/µg DNA of M.CviPI GpC methyltransferase (in-house production) supplemented with 1.2 mM SAM (NEB, B9003S), followed by one 30 min incubation at 37°C with 16 U/µg DNA of M.SssI CpG methyltransferase (NEB, M0226L) in the presence of 7.5 µM MgCl2 and 0.8 mM SAM (NEB, B9003S). Reactions were chemically stopped by addition of 14 µM proteinase K and Stop Solution A (20 mM Tris-HCl, 600 mM NaCl, 1% SDS, 10 mM EDTA) and incubated for at least 4 hour at 55°C. DNA was then purified by phenolchloroform extraction followed by glycogen-assisted salt precipitation. For sample 3 (mA-only), gDNA was treated exclusively with the adenine methyltransferase EcoGII (NEB, M0603S) in two consecutive 30 min incubations at 37°C with 25 U/µg DNA of EcoGII each, supplemented with 1.2 mM SAM, and omitting CpG and GpC methyltransferases. For sample 7 (GpC/CpG/mA), gDNA was treated with M.CviPI as for sample 4 (two consecutive 30 min incubations at 37°C, 8 U/µg DNA M.CviPI each), but 25 U/µg DNA of EcoGII was also added to each incubation. Next, gDNA was treated with an additional 30 min incubation with 16 U/µg DNA of M.SssI in the presence of of 7.5 µM MgCl2 and 0.8 mM SAM as for sample 4. All reactions were terminated and purified as described for sample 4. Unmethylated (sample 0) and fully methylated (samples 3, 4, and 7) gDNA libraries were prepared using the ONT V14 multiplex ligation kit (SQK-NBD114.24) following manufacturer’s instructions. Details on the protocol for the different samples are summarised in Table S4.

### Data alignment

*Drosophila melanogaster* data were aligned against the `BSgenome.Dmelanogaster.UCSC.dm6` genome.

*Mus musculus* data were aligned against the `BSgenome.Mmusculus.UCSC.mm10` genome.

### Short-read sequencing data pre-processing

For short-read datasets (ChIP-seq, DNase-seq, and RNA-seq), reads were aligned using QuasR (60) against the *Drosophila* melanogaster genome (`BSgenome.Dmelanogaster.UCSC.dm6`), or mouse genome (`BSgenome.Mmusculus.UCSC.mm10`).

Publicly available ChIP-seq datasets were pre-processed using the nf-core pipeline nf-core/chipseq v.2.0.0 with arguments `-genome dm6 -macs_gsize 145000000 -aligner star` (61). For our ChromSMF benchmarking against ChIP-seq, we calculated the coverage within each 500 bp tile using the qCount function of the QuasR package (60) with default parameters. We then used this per-tile coverage, to benchmark ChromSMF marked fraction frequencies (Figures 2A, S3E-I). Tiles falling into a blacklisted region as define by the ENCODE project’s datasets (`ENCODE_dm6-blacklist.v2.bed`) were excluded from the analysis.

For our H3K4me3-positive definition, we used a threshold of 100 counts. Tiles with H3K4me3 qCounts > 100 were classified as H3K4me3-positive (4963 tiles), they were classified as H3K4me3-negative otherwise (10904 tiles) (Figure S2C). H3K4me3-positive tiles were further split into four equally populated quantiles (q1-q4, 1241 tiles each) depending on their qCount value. BigWig tracks were generated with the bamCoverage v3.5.0 tool from deepTools with the following command:

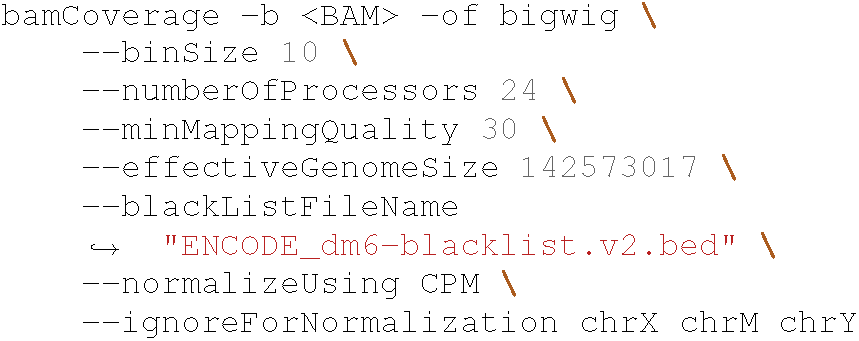

Publicly available DNase-seq datasets were processed as previously described (23).

### Nanopore sequencing data pre-processing

Nanopore data were processed using the following Nextflow pipeline (23): https://git.embl.de/grp-krebs/nf-smfont. The processed methyl BED (mBED) and SMF bulk BigWig files were generated by the pipeline. For detailed and updated description, please refer to the repository.

The unaligned BAM files (uBAM) were generated using Picard RevertSam v3.1.0 (62) to remove alignment information from the BAM files generated by the pipeline. With the following command line:

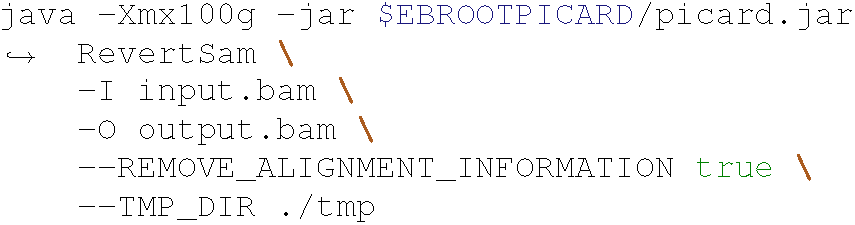

### Basecalling and methylcalling from ONT data

Basecalling, methylation calling, and genomic alignment were performed using Bonito v0.7.3 (with integrated Minimap2 v2.24). We used Bonito since it is the only base-caller that supports methylation calling for cytosines in two specific dinucleotide contexts (CpG and GpC), to date. The super accuracy model v4.2.0 (dna_r10.4.1_e8.2_400bps_sup@v4.2.0) was used for basecalling. All reads with a q-score below 7 were removed from the data. To call cytosine methylation, previously published methylation model trained on CpG and GpC contexts was used (23). To call adenine methylation, the custom-trained methylation model generated with this study was used (See Methods). One BAM file per modification type (mC and mA) was generated. Methylation information was stored in the MM and ML tags. For each BAM file, methylation signals were extracted using pysam, and converted into mBED format (as described in https://github.com/timplab/nanopore-methylation-utilities). The resulting mBED files were indexed using tabix (HTSlib v1.19.2). Methylation probabilities were transformed into log-likelihood ratios (LLRs) for retro-compatibility with previous methylation callers. A stringent absolute LLR cutoff of 2 (corresponding to a p-value of approximately 0.12) was used to classify cytosines or adenines as methylated or unmethylated. Cytosines and adenines for which the model assigned uncertain predictions (−2 > LLR < 2) were excluded from downstream analyses.

### Training and validation of the methyl-caller model for mA

For each sample, Nanopore raw data were basecalled using dorado base-caller 0.3.4 using the base call model dna_r10.4.1_e8.1_400bps_sup@4.2.0 and mapped on the reference *Drosophila* genome (dm6). All reads with a q-score below 7 were removed from the data. Each dataset was split into two uneven groups: 70% of the reads were used for training and 30% for testing model accuracy. The methylation calling model was trained using remora v3.2.0 in a convolutional neural network framework. In summary, the training datasets was generated using the raw signal (pod5 file) and the basecalled data (BAM file) with the following syntax:

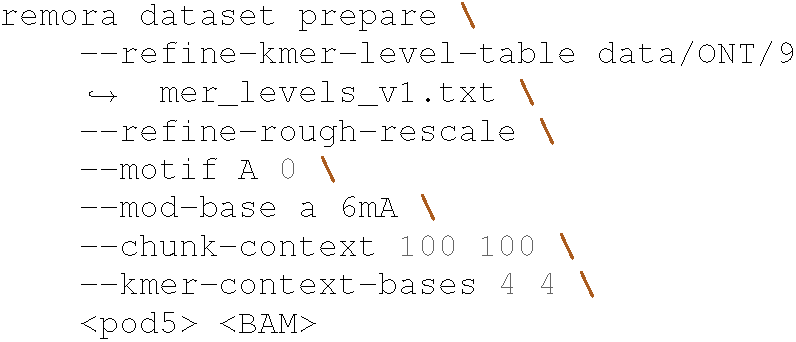

This generated 12,726,282 chunks from 870,023 reads for the fully unmethylated sample, 10,054,515 chunks from 688,673 reads for the mC only, 7,504,665 chunks from 513,428 reads for the mA only, 3,935,490 chunks from 269,203 reads for the fully methylated one. The training dataset was created by taking 70% of each dataset and merging all four ground truth datasets with equivalent weight. The model was then trained in 10 consecutive epoch using the command:

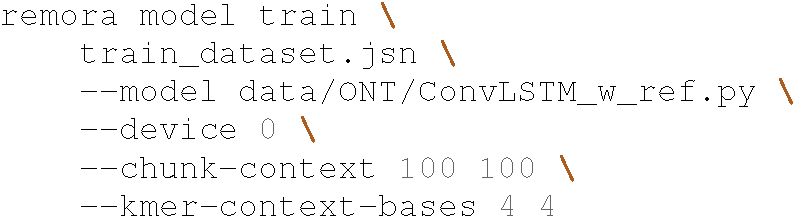

To test the model accuracy, adenine methylation status was inferred using the `remora infer from_pod5_and_bam` tool. The confusion matrix was computed using the `remora validate from_modbams` tool and the results (Figure S1C) plotted using a custom R script. Our custom methylation calling 5 kHz model corrects systematic errors observed in models trained solely on mA provided by ONT (Figure S1B).

### Genome Tiling

Genome information from the `BSgenome.Dmelanogaster.UCSC.dm6` package was used. We kept only standard chromosomes and tiled the *Drosophila* genome in 275’139 non-overlapping tiles using the following command in R: `tileGenome(tilewidth=500, cut.last.tile.in.chrom=TRUE)`

### TSS definition

TSSs for both *Drosophila* and Mouse genomes were defined as previously described (63). Only one TSS per gene was kept.

### Promoter definition

We defined promoters as regions with intermediate CA as detected by the methyLasso software (64) that do overlap at least one TSS. For each promoter, we classified molecules according to their HM status for H3K4me3, we quantified CA separately for HM-positive and HM-negative subpopulations, and we tested for accessibility differences between these two.

### Enhancer definition

We defined enhancers as regions with intermediate CA as detected by the methyLasso software (64) that do not overlap any TSS (12112 enhancers). For each enhancer, we classified molecules according to their HM status for H3K27ac, we quantified CA separately for HM-positive and HM-negative subpopulations, and we tested for accessibility differences between these two.

### Single-molecule methylation frequencies

Singlemolecule methylation state calling for all cytosines and adenines was performed using the nf-smfont pipeline as previously described (23). Reads with per-read mC below 5% and above 95% were filtered out to avoid biases due to DNA coming from incomplete cell permeabilization (0% mC) or dead cells (100% mC).

### Single Nucleotide aggregated methylation frequencies

Single nulceotide methylation frequencies were calculated for each GpC, CpG, or A with at least 20-fold coverage. To perform genome-wide comparison of different samples, mC was directly compared for each GpC and/or CpG (Figures S1E-F, S1I, S3A), while mA was first averaged over 150 bp non-overlapping tiles and then compared (Figures S1G-H, S3B).

### Single-molecule classification for Histone Modifications

Single-molecule classification was performed using the custom `extract_SMinfo_from_mBED.r` script. We integrated the methylation state of adjacent adenines to robustly separate marked molecules from background methylation (Figure S2A-B). For each molecule we extracted adenine methylation rate in the 500 bp tile (see Methods). Only molecules spanning the entire window and having at least 50 informative adenines were considered. Average mA was computed across all informative adenines in the window. Molecules with average mA in the window > 10% were classified as marked (“1”), or classified as unmarked (“0”) otherwise. For each tile, the fraction of marked molecules was computed as the number of molecules classified as marked, over the total number of molecules spanning the extraction window. Only tiles with more than 20 reads spanning it were considered for downstream analysis. To define the fraction of marked molecules at cREs (*i*.*e*., TSSs and enhancers), we used the single-molecule classification information of the tile they fall into as a proxy for the cRE status.

### Single-molecule classification for Nucleosome Phasing at TSSs

Single-molecule classification was performed using the nf-smfont pipeline (23). For each molecule we extracted cytosine methylation rate in a 101 bp window centred at +135 bp downstream the TSS, as this was previously described to be the position of phased nucleosome during transcription in *Drosophila* (65). Only molecules spanning the entire window were considered. Average mC was computed across all cytosines in the window. Molecules with average mC in the window > 50% were classified as accessible (“1”), or classified as closed (“0”) otherwise. Nucleosome phasing was defined as molecule being closed (“0”) in the window. For each TSS, the fraction of accessible molecules was computed as the number of molecules classified as accessible, over the total number of molecules spanning the TSS extraction window. Only TSSs with more than 20 reads spanning it were considered for downstream analysis.

### Single-molecule classification for Chromatin Accessibility at enhancers

Single-molecule classification was performed using the nf-smfont pipeline (23). For each molecule we extracted cytosine methylation rate in a 101 bp window centred on the enhancers as defined by the methyLasso analysis (see Methods). Only molecules spanning the entire window were considered. Average mC was computed across all cytosines in the window. Molecules with average mC in the window > 50% were classified as accessible (“1”), or classified as closed (“0”) otherwise. For each enhancer, the fraction of accessible molecules was computed as the number of molecules classified as accessible, over the total number of molecules spanning the enhancer extraction window. Only enhancers with more than 20 reads spanning it were considered for downstream analysis.

### Comparison of ChromSMF single-molecule quantification with ChIP-seq counts for H3K4me3 at TSSs

For each 500 bp tile across the genome, we computed the fraction of molecules classified as H3K4me3-marked at the tile. We then restricted our analysis to tiles spanning at least one TSS, and compared these frequencies with H3K4me3 counts (log2) as measured by ChIP-seq at the same 500 bp tiles. We added a pseudocount of 10 to ChIP-seq counts to avoid issues with log transformation (Figure 2A). Tiles falling into a blacklisted region as define by the ENCODE project’s datasets (`ENCODE_dm6-blacklist.v2.bed`) have been excluded from the analysis (7621 tiles, 2.8%). Tiles at sex chromosomes and at mitochondrial DNA have been excluded from the analysis (8246 tiles, 3.0%). Pearson’s correlation coefficient was computed using the cor function from base R with default parameters. Correlations were calculated on log2 transformed ChIP-seq data.

### Quantification of the association between H3K4me3 and nucleosome phasing at TSSs

Single-molecule classification at each TSS was performed independently for H3K4me3 and nucleosome phasing (see Methods). Information was then crossed using the custom `smf_sort_extract_dimelo.R` script, which relies on consistent readIDs between mC and mA analysis. For each TSS, the fraction of accessible molecules for the two HM fractions (H3K4me3-marked and unmarked) was computed. Delta in nucleosome phasing was computed as the difference between the fraction of accessible molecules in the marked fraction - fraction of accessible molecules in the unmarked fraction. DeltaCA < 0 indicates H3K4me3-marked molecules have higher nucleosome positioning at the TSS. Two-sided Fisher’s exact test was performed to test for significance in nucleosome positioning differences. TSSs with adjusted p-value <= 0.05 and log2OR <= −1 were classified as significant.

### Quantification of the association between H3K27ac and chromatin accessibility at enhancers

Singlemolecule classification at each enhancer was performed independently for H3K27ac and chromatin accessibility (see Methods). Information was then crossed using the custom `smf_sort_extract_dimelo.R` script, which relies on consistent readIDs between mC and mA analysis. For each enhancer, the fraction of accessible molecules for the two HM fractions (H3K27ac-marked and unmarked) was computed. Delta in accessibility was computed as the difference between the fraction of accessible molecules in the marked fraction - fraction of accessible molecules in the unmarked fraction. DeltaCA > 0 indicates H3K27ac-marked molecules have higher accessibility at the enhancer. Two-sided Fisher’s exact test was performed to test for significance in accessibility differences. Enhancers with adjusted p-value <= 0.05 and log2OR >= 1 were classified as significant.

### Quantification of long distance association

Single-molecule classification at each cRE was performed independently for HM and CA (see Methods). Information was then crossed using the custom `smf_sort_extract_dimelo.R` script, which relies on consistent readIDs between mC and mA analysis. For each cRE of interest (*i*.*e*., enhancer or promoter) we classified molecules based on the presence, or absence of the HM (*i*.*e*., H3K27ac or H3K4me3, respectively) at that cREs (mA sorting viewpoint) and computed delta in CA between the two fractions at the same cRE (mA sorting viewpoint). Two-sided Fisher’s exact test was performed to test for significance in accessibility differences. Only cREs (mA sorting viewpoint) with adjusted p-value <= 0.05 and log2OR >= 1 were classified as significant and considered for the analysis.

We then evaluated differences in CA between the HM-marked and unmarked molecules. We looked at CA on the same molecules but at the nearby cREs (mC sorting distal cRE). Considering the average read length of our datasets, only cREs falling within +-5 kb from the sorting viewpoint were considered in the analysis. Delta in accessibility was computed as the difference between the fraction of accessible molecules (at the mC sorting distal cRE) in the HM-marked fraction - fraction of accessible molecules in the unmarked fraction (at the mA sorting viewpoint). DeltaCA > 0 indicates HM-marked molecules at the viewpoint have higher accessibility at the distal cRE. Two-sided Fisher’s exact test was performed to test for significance in accessibility differences. Distal cREs with adjusted p-value <= 0.05 and log2OR >= 1 were classified as significant.

### Alleles separation based on known phased genome

SNPs between Bl6 and CAST genomes were retrieved as a VCF file from the EBI dataset (https://ftp.ebi.ac.uk/pub/databases/mousegenomes/REL-1505-SNPs_Indels/). We kept only variants with the FILTER=PASS status, filtering out variants which are heterozygous and/or with low confidence calling. Since the genotype in this file is relative to the full CAST species, we then manually set all the variants to heterozygous. We then generated a phased VCF which would represent the F1 genome (*i*.*e*., all variants are 0|1).

Haplotype assignment was performed on mC-BAM file using WhatsHap haplotag (v.2.6, (56)) and the following parameters:

`-reference mm10.fa -ignore-read-groups`

`-skip-missing-contigs -tag-supplementary`.

Then, readIDs were used to also tag the mA-BAM file.

### Alleles separation inferred from the Nanopore data

SNPs between Bl6 and CAST genome were retrieved as a VCF file from the EBI dataset: https://ftp.ebi.ac.uk/pub/databases/mousegenomes/REL-1505-SNPs_Indels/.

VCF file was then genotyped and phased using WhatsHap (v.2.6, (56)) with the following parameters:

`-ignore-read-groups -reference mM10.fa`.

Haplotype assignment was performed on mC-BAM file using WhatsHap haplotag (v.2.6, (56)) and the following parameters:

`-reference mM10.fa -ignore-read-groups`

`-skip-missing-contigs -tag-supplementary`. Then, readIDs were used to also tag the mA-BAM file.

### ICR definition

ICR were defined as previously described (8). TSS were assigned to ICRs using the subsetByOverlap function. ICRs with no differences in 5mC between haplotypes were excluded from the analysis.

### Extraction and analysis of mC for F1 mouse data with endogenous 5mC

Single-molecule methylation state calling for all cytosines with at least 10-fold coverage was performed as described above. Yet, some contexts were excluded from the analysis as previously described (8) to avoid interferences between CpG and GpC methylation calling. For chromatin accessibility analysis, only cytosines in D**GC**HN (meaning [AGT]**GC**[ACT][ACGT]) contexts were considered. For endogenous DNA methylation analysis, only cytosines in NW**CG**W (meaning [ACGT][AT]**CG**[AT]) contexts were considered.

### Haplotype resolved analysis

For each TSS, we extracted single-molecule information separately for each channel (*i*.*e*. mA for H3K4me3, mC at CpGs for 5mC, and mC at GpCs for CA), and then computed frequencies for either combined or individual alleles.

We restricted our analysis to one TSS per gene and to only the ones that do not have other TSSs within +-1 kb window. For both filtering, only the most 5’ TSS was retained.

We classified single molecules depending on their H3K4me3 status at those TSSs. We extracted the mA fraction for each read at 100 bp window around the +1 liker DNA, resulting window is [+165;265] bp from the TSS. Only molecules with at least 10 informative adenines were considered. Molecules with mA > 10% were classified as H3K4e3-marked at that promoter, H3K4me3-unmarked otherwise. Per-haplotype H3K4me3-marked frequencies (SM-freq_HapX) were computed back using total counts and haplotagged readIDs.

Given the low coverage we used average 5mC and CA at each TSS as a proxy for single-molecule measurements. This is acceptable as we don’t perform single-molecule association analysis with this datasets, but we only analysed the global average signal for each haplotype.

For each TSS of the analysis, we computed the average mC for each cytosine at CpGs within [-1000;+200] from the TSS. Only TSSs with at least 5 cytosines were kept for the analysis. If the average-average mC was > 50% we classified the TSS as methylated. Per-haplotype 5mC averages (avg5mC_HapX) were computed the same way but using haplotype-specific frequencies of mCpG.

For each TSS of the analysis, we computed the average mC for each cytosine at GpCs within [-100;+50] from the TSS. Only TSSs with at least 5 cytosines were kept for the analysis. If the average-average mC was > 50% we classified the TSS as accessible. Per-haplotype chromatin accessibility (avgCA_HapX) were computed the same way but using haplotype-specific frequencies of mGpC.

Delta for each feature were computed as follows. For 5mC: `avg5mC_Hap1 - avg5mC_Hap2`; for CA: `avgCA_Hap1 - avgCA_Hap2`; for HM: `SMfreq_Hap1 - SMfreq_Hap2`.

### RNA-seq analysis and identification of genes with allelic imbalance

VCF file was either the manually curated one or the one generated from nanopore data depending on the analysis.

Deseq2 analysis was done Hap1/Hap2. Fold change in gene expression was computed. Genes with abs(expr_log2FC) >= 1 and expr_padj < 0.05 were classified as imbalanced.

### Statistical analysis and reproducibility

All statistical analyses were performed using R software and the R package tidyverse (66) and are described in the figure legends and methods section. Violin and box plots were plotted by the R package ggplot2 (67) with `outlier.shape = NA and trim = TRUE`. The upper and lower boundaries of the box plot represent the 25th and the 75th percentiles. The central line represents the median. Sample size for each violin is shown when relevant (Figures 4B, S4B-C, and S6B). Pearson’s correlation coefficients in Figures 2A, S1E-I, S2E, and S3A-H were calculated using the R package corr (68) with the option `pairwise.complete.obs`. The twosided Fisher’s exact test (Figures 2E, and 3D) was performed under the criteria that the region of interest was covered by at least 20 reads. ChromSMF single locus plots (Figures 1B-D, 2F, 3E, 4D, 5B-D, 6C, S2F, S3I, S4D, and S5E) were plotted through ggplot2 and custom scripts.

### Scripting, Data Analysis, and High-Performance Computting

All the scripting and data analysis for this project were performed using R-4.2.2 for the nf-smfont pipeline and R-4.4.1 for the downstream analysis. The pre-processing pipeline for SMF and ChromSMF data was scripted using Nextflow-24.04.2 (69). All high-performance computing was performed using the SLURM workload manager (70), either through direct bash scripting or through the R package rslurm (71).

## Acknowledgments

We would like to thank Laura Moniot-Perron, Colm Doyle, Elisa Kreibich, Rob Klose, and Rodrigo Villaseñor for comments on the manuscript. Mathias Boulanger for initial support to develop the computational pipeline. The EMBL PepCore facility for the production of the pA-Hia5 and M.CviPI MTases. The EMBL GeneCore facility for support with P2 ONT sequencing. The DKFZ OpenLab for providing *ad hoc* access to the P24 PromethION instrument. Rozemarijn Kleinendorst for extracting RNA to perform RNAseq for the F1 mESCs. MODIS and Charles Girardeau for support with data management. Nicolas Altemose for technical suggestions, and providing aliquots of the pA-Hia5 construct for initial benchmarking. Tobias Rausch for suggestions on the data-driven reconstruction of haplotypes. Marco Alecci for working on the formatting of the LaTeX version of this paper. Research in the laboratory of A.R.K. is supported by core funding from the EMBL, Deutsche Forschungsgemeinschaft (KR 5247/1-2, KR 5247/1-3) and the ERC (TFCoop-101125530).

## Contributions

A.R.K. designed the study. M.P. and A.R.K. wrote the manuscript. M.P. developed and optimized the ChromSMF protocol, performed the experiments, and analysed the data. M.P. trained the ONT methyl-caller model for mC-insensitive adenine methylation detection. A.R.K supervised conduction of the experiments and the data analysis. All authors discussed the results and commented on the manuscript.

## Additional Information

## Supplementary Figures

**Fig. S1.**
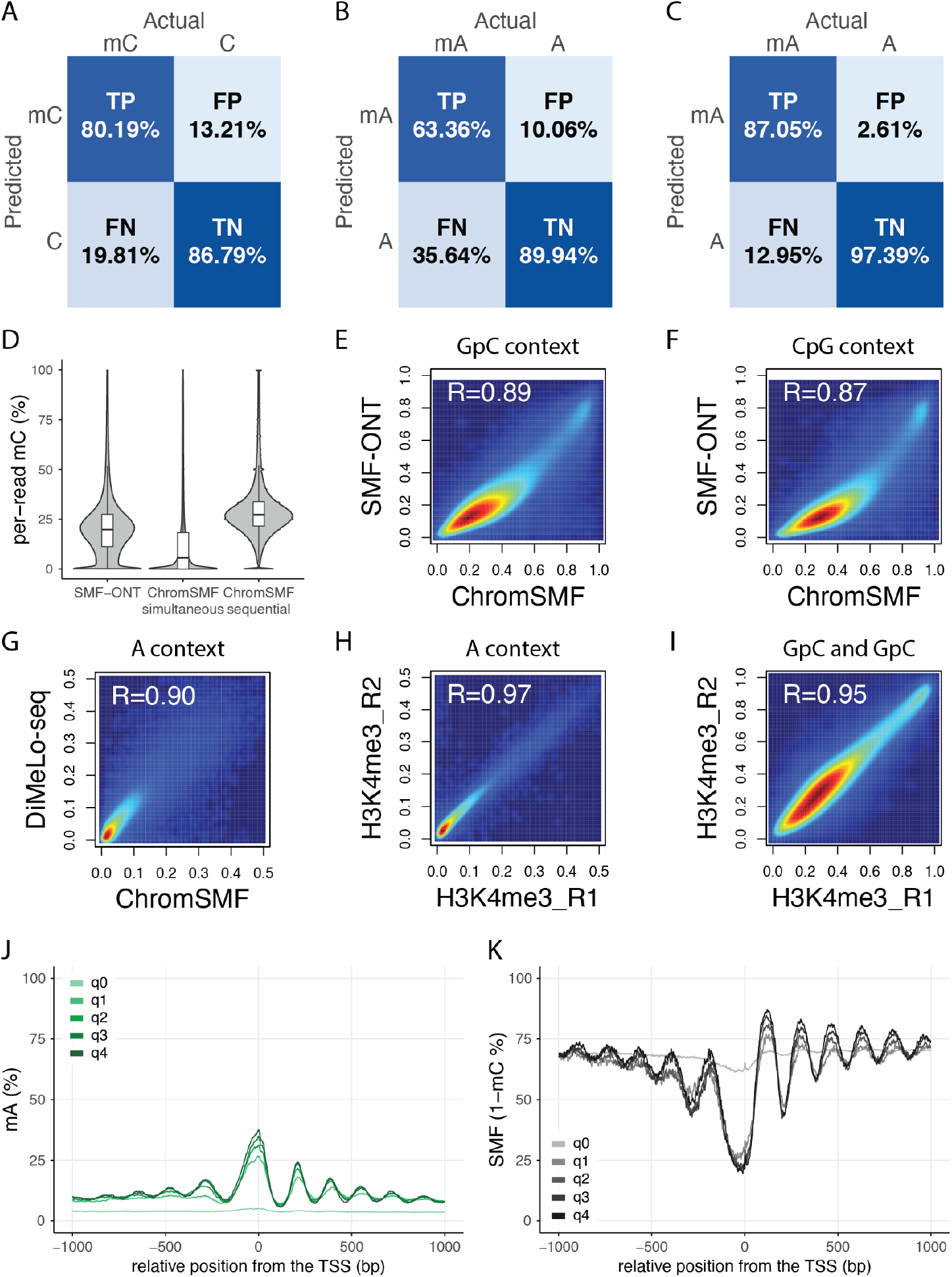
Supplementary Figure relative to Fig.1. **(A) Confusion matrix for mC detection when methylation in A context is present**. The model (23), trained on fully methylated (CpG and GpC dinucleotides only) and fully unmethylated DNA, detects mC with 83.5% accuracy on 5 kHz ONT data. (TP = true positives, TN = true negatives, FP = false positives, FN = false negatives). **(B–C) Confusion matrix for mA detection when methylation in both CpG and GpC contexts are present. (B)** Confusion matrix of the ONT model `dna_r10.4.1_e8.2_5khz_400bps_sup_v4.2.0_6ma_v3.pt`. The model, trained on A and mA data only, detects mA with 77.1% accuracy on 5 kHz ONT data. **(C)** Confusion matrix of the custom convolutional neural network model optimised for ChromSMF data. The model, trained on all possible combinations of methylation in CpG, GpC, and A contexts, detects mA with 92.2% accuracy on 5 kHz ONT data. (TP = true positives, TN = true negatives, FP = false positives, FN = false negatives). **(D) Sequential but not simultaneous ChromSMF methylations recapitulate SMF signal**. Distribution of per-read mC in SMF-ONT, ChromSMF with simultaneous methylation, and ChromSMF with sequential methylations (left to right). **(E–F) ChromSMF correlates with SMF**. Smoothed scatter plots comparing the average cytosine methylation in **(E)** GpC and **(F)** CpG contexts where the mC footprinting treatment was performed individually (SMF-ONT) or in combination with histone modification detection (ChromSMF). *R* represents the Pearson’s correlation coefficient. **(G) ChromSMF correlates with DiMeLo-seq for H3K4me3**. Smoothed scatter plot comparing the average adenine methylation at 150 bp tiles across the genome where mA-based H3K4me3 detection was performed individually (DiMeLo-seq) or in combination with the mC-based footprinting (ChromSMF). *R* represents the Pearson’s correlation coefficient. **(H–I) ChromSMF correlates across replicates for H3K4me3**. Smoothed scatter plots comparing the average methylation at adenines over 150 bp tiles **(H)**, or at single GpCs and CpGs **(I)** across the genome. *R* represents the Pearson’s correlation coefficient. **(J–K) ChromSMF signal scales with H3K4me3 ChIP-seq enrichment**. Composite H3K4me3 (mA%) **(J)** or chromatin accessibility (1 – mC%) **(K)** signal within 2 kb around TSSs genome-wide. TSSs are split into five groups based on ChIP-seq enrichment for H3K4me3: q0 comprises H3K4me3-negative TSSs (ChIP-counts *<* 100), and q1–q4 represent equal-sized quantiles of H3K4me3-positive TSSs (ChIP-counts ≥ 100). Signal is smoothed every 5 nt.

**Fig. S2.**
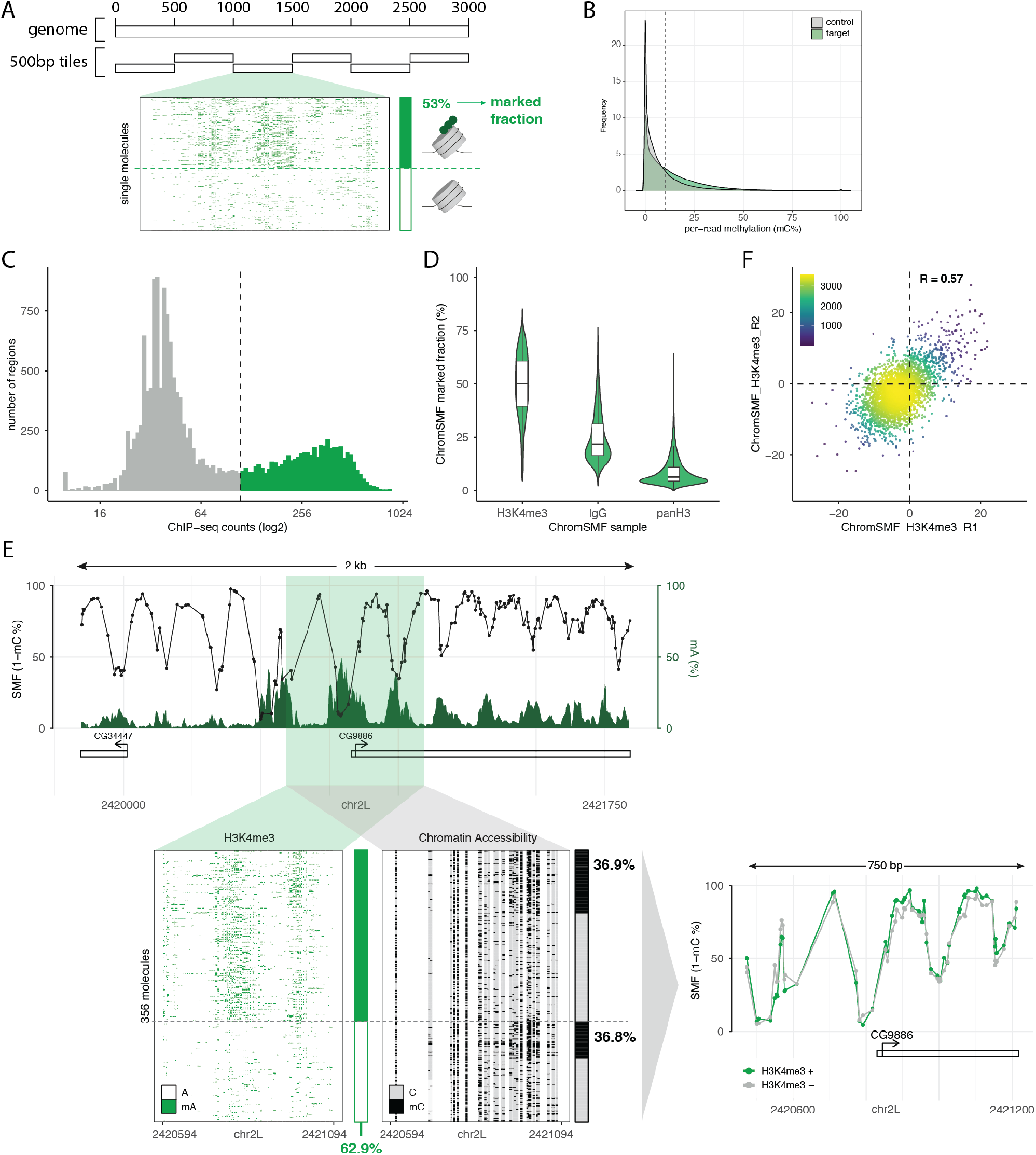
Supplementary Figure relative to Fig.2. **(A) Computational strategy to extract single-molecule frequencies of histone modifications**. The genome is tiled into 500 bp non-overlapping windows. Molecules with an average mA ≥ 10% within a tile are classified as HM-marked at that tile, and unmarked otherwise. Molecules with fewer than 50 informative adenines in the tile are excluded from the analysis. **(B) mA distribution distinguishes H3K4me3-enriched and control regions**. Comparison of single-molecule mA% distributions in the H3K4me3 ChromSMF experiment between tiles enriched for H3K4me3 by ChIP-seq (target regions; green) and negative regions (control regions; grey). The dashed line indicates the adenine methylation threshold (10% mA per-read per-tile) used to classify molecules as marked. **(C) Distribution of H3K4me3 ChIP-seq counts across 500 bp tiles**. Tiles with counts *>* 100 (dashed line) are classified as H3K4me3-positive (green), while the remaining tiles are classified as H3K4me3-negative (grey). ChIP-seq counts are shown on a logarithmic scale (x-axis; log_2_). **(D) ChromSMF mA marking is specific to the antibody used**. Boxplots comparing the fraction of marked molecules at H3K4me3-positive tiles in ChromSMF samples generated with an H3K4me3 antibody, a non-specific IgG control, or a pan-H3 antibody (left to right). **(E) Single locus example of the *CG9886* promoter where no association between H3K4me3 and chromatin accessibility is detected**. Top panel: ChromSMF signal for H3K4me3. Average SMF signal (1 – mC%) of individual cytosines (black) and average smoothed mA signal (mA%; green; smoothing across 4 adenines). Bottom panel - left: single-molecule stacks displaying either mA-H3K4me3 (green, left) or mC-chromatin accessibility (black, right) signal. Molecules are displayed in identical order in both panels and originate from the same sample. Single-molecule classification of H3K4me3 (green) and chromatin accessibility (black) are shown as stacked barplots on the right of each stack. Single-molecule quantification of total H3K4me3 at the locus is shown at the bottom. Single-molecule classification of chromatin accessibility for H3K4me3-marked and H3K4me3-unmarked molecules is shown on the right. Bottom panel - right: average SMF signal (1 – mC%) of individual cytosines for molecules classified as either marked (green) or unmarked (grey) by H3K4me3. **(F) Molecular associations between H3K4me3 and chromatin accessibility are reproducible across replicates**. Smoothed scatter plot comparing delta-delta H3K4me3 abundance between replicates at TSSs. *R* represents the Pearson’s correlation coefficient.

**Fig. S3.**
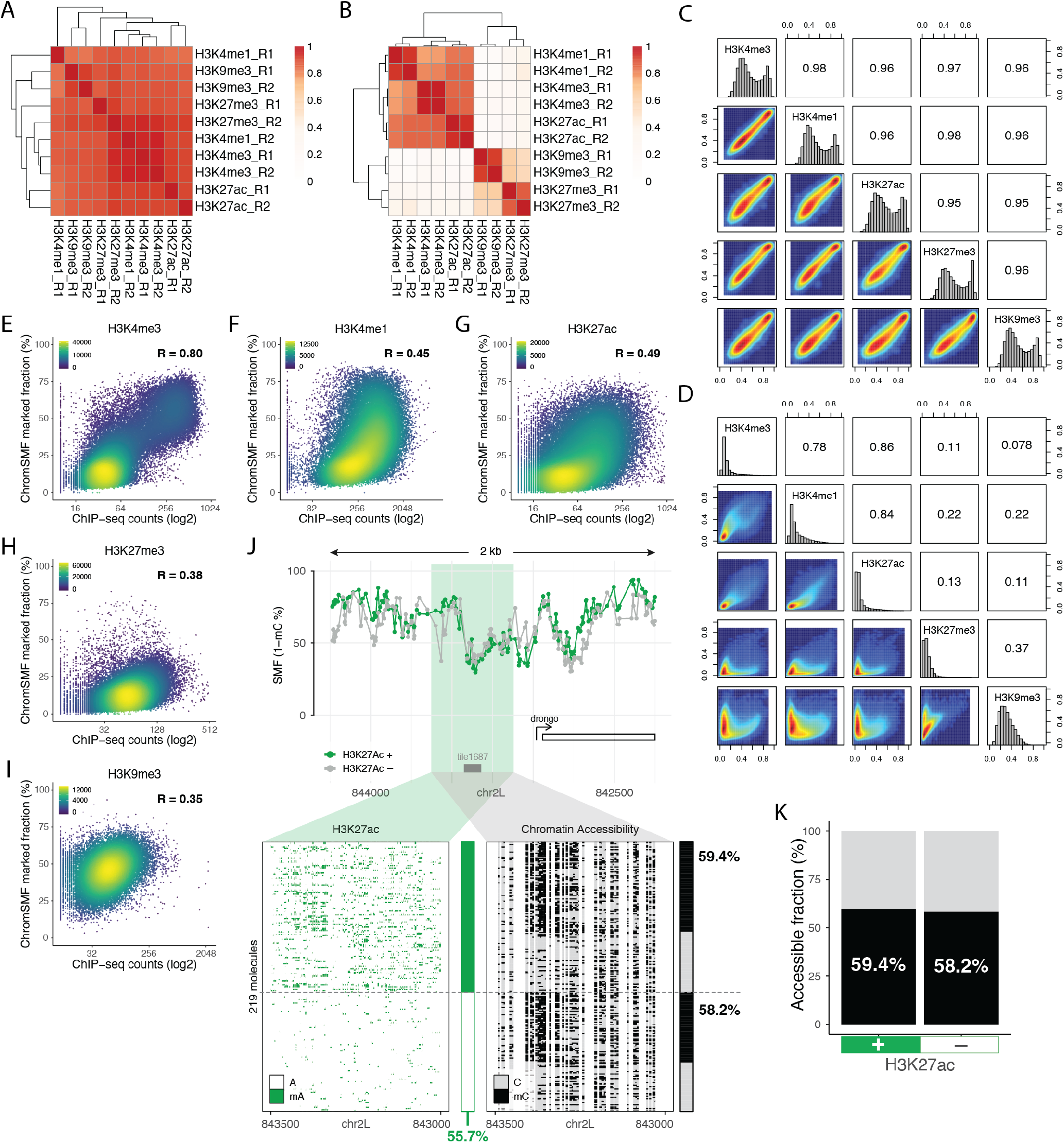
Supplementary Figure relative to Fig.3. **(A) ChromSMF mC signal is comparable across samples**. Pearson correlation matrix of single-cytosine average mC levels (CpG and GpC contexts) in ChromSMF samples. **(B) ChromSMF mA signal is specific to the targeted histone modification and reproducible across replicates**. Pearson correlation matrix of average mA levels across 150 bp genomic tiles in ChromSMF samples. **(C) Single-molecule quantification of chromatin accessibility is comparable between samples**. Pairwise smoothed scatter plots of the fraction of accessible molecules at DHSs for ChromSMF experiments targeting different histone modifications. **(D) Single-molecule quantification of histone modifications recapitulates known chromatin relationships**. Pairwise smoothed scatter plots of the fraction of molecules classified as marked by each histone modification in different ChromSMF experiments. **(E–I) Benchmarking of the single-molecule quantification of histone modifications**. Scatter plots comparing the fraction of molecules marked by a specific histone modification as defined by ChromSMF (y-axis) against the ChIP-seq counts (log_2_) for the same modification at the same genomic tile (x-axis). Data were collected from 500 bp tiles genome-wide and annotated using ChromHMM data. *R* represents the Pearson’s correlation coefficient. Data are presented for **(E)** H3K4me3 at TSSs, gene bodies (transcription elongation), and open chromatin regions; **(F)** H3K4me1 at enhancers and open chromatin regions; **(G)** H3K27ac at enhancers, open chromatin, and TSSs; **(H)** H3K27me3 at Polycomb-mediated repressed regions and transcriptionally silent intergenic regions; and **(I)** H3K9me3 at pericentromeric heterochromatin and heterochromatin-like regions embedded within euchromatin. **(J–K) Single locus example of a locus where no molecular association between H3K27ac and chromatin accessibility is detected. (J)** ChromSMF signal for H3K27ac. Top panel: average SMF signal (1 – mC%) of individual cytosines for molecules classified as either marked (green) or unmarked (grey) by H3K27ac. Bottom panel: single-molecule stacks displaying either mA-H3K27ac (green, left) or mC-chromatin accessibility (black, right) signal. Molecules are displayed in identical order in both panels and originate from the same sample. Single-molecule classification of H3K27ac (green) and chromatin accessibility (black) are shown as stacked bar plots on the right of each stack. Single-molecule quantification of total H3K27ac at the locus is shown at the bottom. Single-molecule classification of chromatin accessibility for H3K27ac-marked and H3K27ac-unmarked molecules is shown on the right. **(K)** Bar plot depicting the single-molecule quantification of chromatin accessibility at the enhancer upstream of the *drongo* promoter for molecules classified as marked (left) or unmarked (right) by H3K27ac at the same enhancer.

**Fig. S4.**
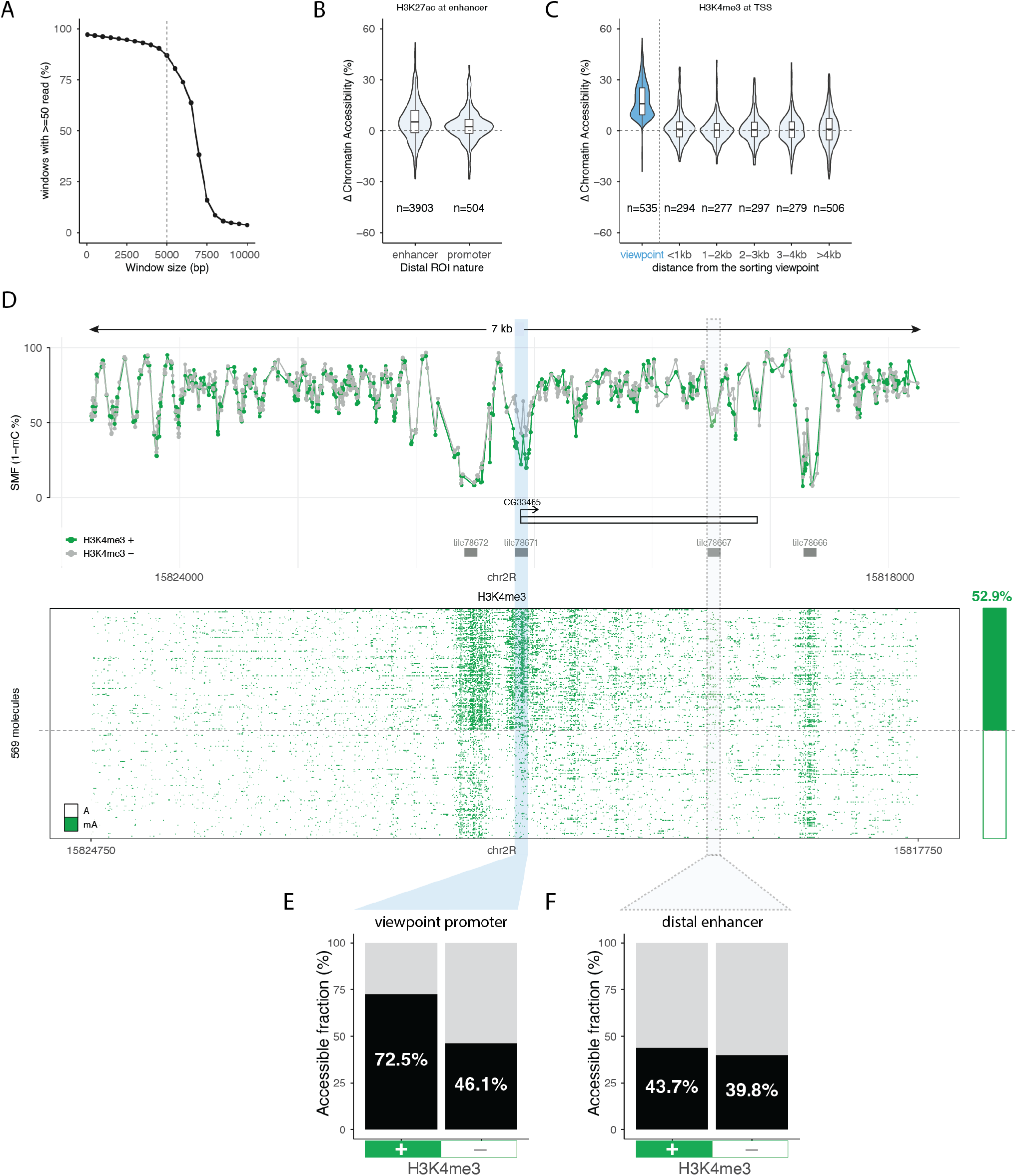
Supplementary Figure relative to Fig.4. **(A) Fraction of windows with more than 50 reads decreases as a function of window size**. The *Drosophila* genome was chunked into non-overlapping windows and the fraction of reads spanning each window was computed. The analysis was repeated across multiple iterations using different window sizes (from 1 bp to 10 kb). For each iteration, the fraction of windows with at least 50 reads was calculated. Example shown for the H3K4me3_R1 sample (average genome-wide coverage 530*×*, average read length 6.1 kb). **(B) The molecular association between H3K27ac and chromatin accessibility at distant loci is observed at both promoters and enhancers**. Distribution of the difference in chromatin accessibility at distal cREs between H3K27ac-marked and unmarked fractions as a function of the nature of the distal cRE. **(C) The molecular association between H3K4me3 and chromatin accessibility is not observed for distant cREs**. Distribution of the difference in chromatin accessibility between H3K4me3-marked and unmarked fractions at the viewpoint (promoters; blue) and at cREs located at various distances from it (white). **(D) Single locus example where H3K4me3 at the *CG33465* promoter is not associated with higher chromatin accessibility at other cREs of the *cis*-regulatory landscape**. Top panel: ChromSMF signal for H3K4me3. Average SMF signal (1 – mC%) of individual cytosines from molecules classified as either marked (green) or unmarked (grey) by H3K4me3. Bottom panel: single-molecule stacks displaying mA-H3K4me3 (green) signal. Single-molecule classification of H3K4me3 at the viewpoint promoter is shown as a stacked barplot on the right. Single-molecule quantification of total H3K4me3 abundance at the viewpoint promoter (green) is shown on top of the barplot. **(E–F) H3K4me3 associates with different chromatin accessibility only at the marked promoter**. Bar plot depicting the single-molecule quantification of chromatin accessibility for molecules classified as H3K4me3-marked (left) or unmarked (right) at the *CG33465* promoter. CA is quantified at the *CG33465* promoter itself **(E)** or at the downstream enhancer **(F)**.

**Fig. S5.**
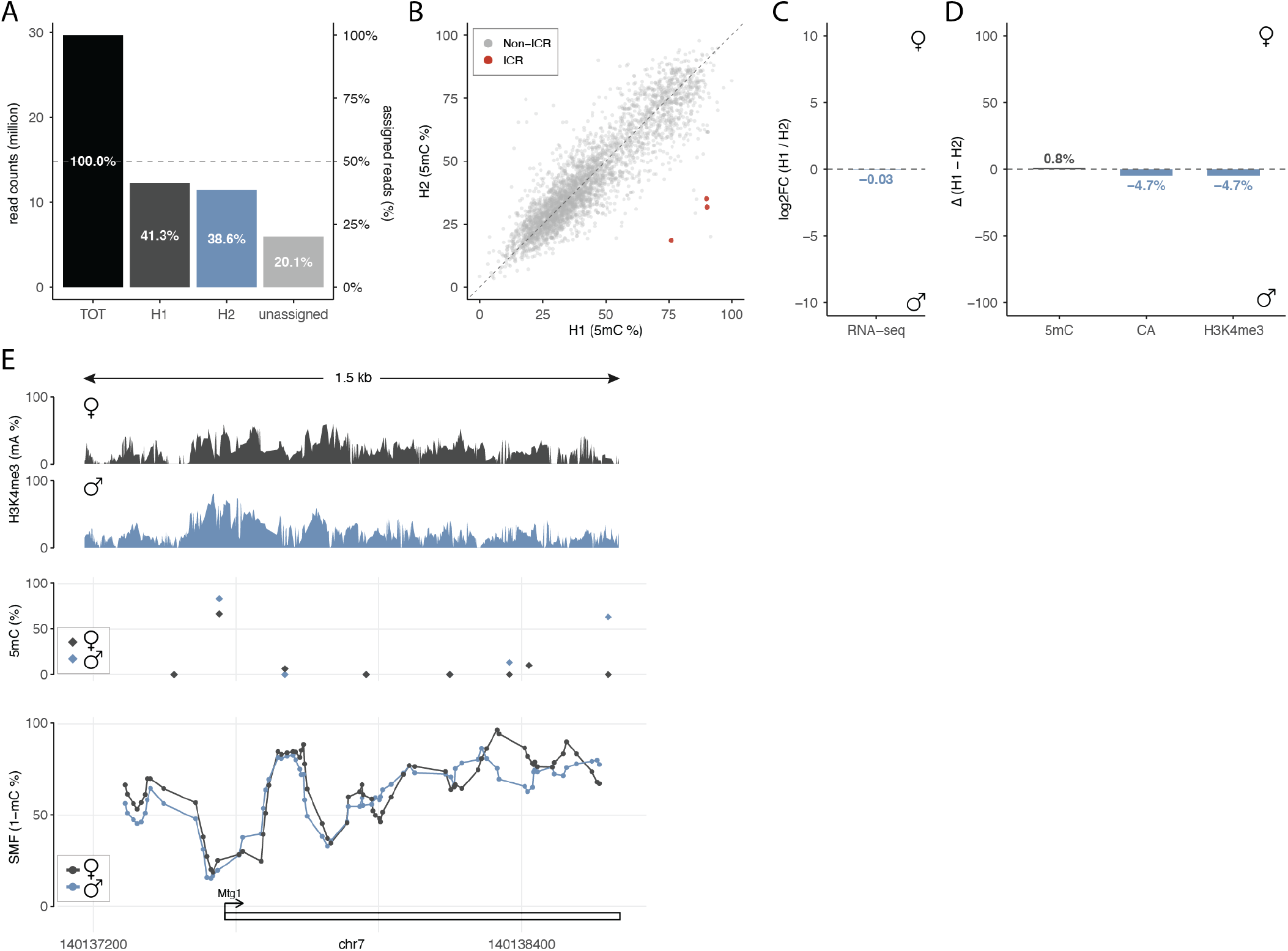
Supplementary Figure relative to Fig.5. **(A) Reads assignment is balanced between alleles**. Bar plot showing the percentage of assigned reads over the total. Reads with haplotype assigned are distributed equally between alleles. Only 20% of the total reads could not be assigned to either allele. **(B) ChromSMF detects differential methylation of ICRs between alleles**. Scatter plot comparing average 5mC at TSSs (mC% in CpG context) between alleles genome-wide. ICRs are highlighted in red. **(C–D) Quantification of allelic similarities at the non-ICR *Mtg1* promoter**. ChromSMF was performed in F1 hybrid (Bl6/CAST) mESCs targeting H3K4me3. **(C)** Bar plot showing log_2_ FoldChange in gene expression between the two haplotypes at the *Mtg1* promoter. **(D)** Bar plot showing haplotype symmetry at the *Mtg1* promoter for endogenous methylation (5mC; average mC% in CpG context), chromatin accessibility (CA; average mC% in GpC context), and H3K4me3 (single-molecule sorting for H3K4me3-mA using a 100 bp window around the TSS). Colours depict in which allele the value is higher (maternal = dark-grey, paternal = blue). Alleles have comparable levels of gene expression, 5mC, CA, and H3K4me3. **(E) Single locus example of allelic similarities at the non-ICR *Mtg1* promoter**. Signals are plotted for the two alleles separately and the colour code reflects haplotype origin: Haplotype1 (H1, ♀, maternal) is represented in dark-grey, Haplotype2 (H2, ♂, paternal) is represented in blue. All signals originate from the same sample. Top panel: Average mA signal (mA%) smoothed every 3 adenines within a 1.5 kb window around the annotated CpG island. Middle panel: Average 5mC signal (mC%) at individual CpGs for the same region. Bottom panel: Average SMF signal (1 – mC%) at the same region, smoothed every 4 GpCs to compensate for low coverage.

**Fig. S6.**
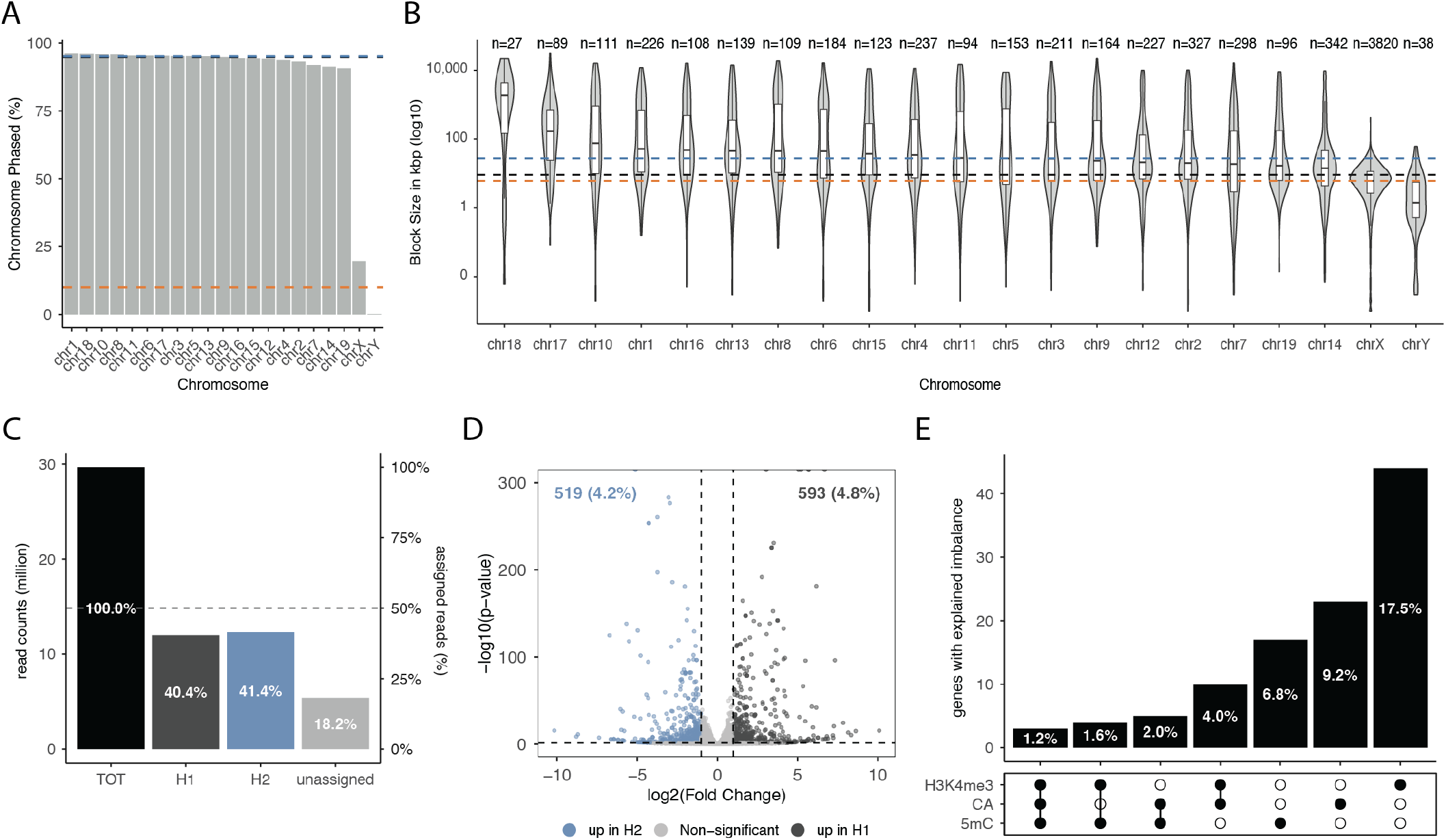
Supplementary Figure relative to Fig.6. **(A) Data-driven reconstruction of haplotypes allows phasing of almost all autosomes**. Bar plot showing the percentage of phased genome per chromosome. Chromosomes are sorted by phased percentage value. Median fraction of phased chromosome for autosomes (blue = 95%), sex chromosomes (orange = 10%), and all chromosomes (black = 94.8%) are shown. **(B) Median phase-block size in autosomes is 28 kb**. Distribution of phase-block size per chromosome. Chromosomes are sorted by median phase-block size value. Median phase-block sizes for autosomes (blue = 27.69 kb), sex chromosomes (orange = 6.12 kb), and all chromosomes (black = 9.08 kb) are shown. **(C) Read assignment using data-driven reconstruction is balanced between haplotypes**. Bar plot showing the percentage of assigned reads over the total. Reads with haplotype assignment are distributed equally between alleles. Only 18% of the total reads could not be assigned to either allele. **(D) Identification of genes with allelic imbalance in gene expression**. Volcano plot depicting the fold change (log_2_ FC) and the p-value (− log_10_pValue) from the DESeq2 analysis testing for differential gene expression between haplotypes. Most genes show no differential gene expression between alleles (grey dots), yet 1112 genes show imbalanced gene expression for either of the two alleles (haplotype 1 = dark-grey, haplotype 2 = blue). **(E) Fraction of genes with allelic imbalance associated with asymmetric epigenetic features**. Genes were classified based on whether allelic differences in 5mC, chromatin accessibility (CA), and H3K4me3 exceeded a threshold of 15%. Each bar represents the number of genes showing a specific combination of epigenetic asymmetries. The dot matrix below the bars indicates the presence (filled circle) or absence (open circle) of haplotype asymmetry for each feature (rows: H3K4me3, CA, 5mC), with vertical connectors highlighting the combination present in each category. Genes with no detectable asymmetry in any feature were excluded from the analysis (145 genes, 58%). Bar labels indicate the fraction relative to all genes with allelic expression imbalance (251 genes).

## Supplementary Tables

**Table S1.** ChromSMF datasets generated in *Drosophila* S2 cells. ChromSMF experiments were performed in *Drosophila* Schneider’s S2 cells. At least two replicates for each targeted histone modification (HMs) were generated. Targeted histone modifications are listed with the type of region where they are predominantly deposited. Highlighted in green are the histone modifications associated with gene activation, in red the ones associated with repression. Three control experiments were performed using a non-specific IgG antibody or a non-modified pan H3 one. Sequencing outputs are shown as per-nucleotide genome-wide coverage (median) and average read length (N50).

| *Drosophila* S2 cells |  |  |  |  |
| --- | --- | --- | --- | --- |
| Modification | Regulatory region | Replicate | Median GW coverage | Average Read Length (N50) |
| **H3K4me3** | promoter (P) | R1 | 530X | 6.1 kb |
|  |  | R2 | 580X | 5.8 kb |
| **H3K4me1** | enhancer (E) | R1 | 150X | 5.8 kb |
|  |  | R2 | 620X | 5.6 kb |
| **H3K27ac** | E & P | R1 | 200X | 4.6 kb |
|  |  | R2 | 200X | 5.2 kb |
| **H3K27me3** | polycomb | R1 | 230X | 5.4 kb |
|  |  | R2 | 710X | 6.2 kb |
| **H3K9me3** | heterochromatin | R1 | 250X | 4.0 kb |
|  |  | R2 | 270X | 5.1 kb |
| **panH3** | non-specific | R1 | 290X | 5.9 kb |
|  |  | R2 | 190X | 5.2 kb |
| **IgG isotype** | non-specific | R1 | 105X | 5.2 kb |

**Table S2.** ChromSMF datasets generated in F1 mESCs (BL6/CAST). ChromSMF experiments were performed in BL6/CAST F1 mouse embryonic stem cells. Replicates for the targeted histone modification H3K4me3 are listed together with sequencing output metrics including per-nucleotide genome-wide coverage (median) and average read length (N50).

| F1 mESCs (BL6/CAST) |  |  |  |  |
| --- | --- | --- | --- | --- |
| Modification | Regulatory region | Replicate | Median GW coverage | Average Read Length (N50) |
| **H3K4me3** | promoter (P) | R1 | 5X (+ 10X AS) | 4.6 kb |
|  |  | R2 | 14X | 6.3 kb |
|  |  | R3 | 6X | 5.0 kb |

**Table S3.** Antibodies used for ChromSMF experiments. ChromSMF experiments were performed in two different cell lines (*Drosophila* Schneider’s S2 cells and BL6/CAST F1 mouse embryonic stem cells). The table lists the antibodies used for each target histone modification, including the commercial source, catalog identifier, and dilution ratio used in the experiments.

| Target | Source | Identifier | Ratio used |
| --- | --- | --- | --- |
| **H3K4me3** | abcam | ab8580 | 1:50 (4 µg) |
| **H3K4me1** | CST | 5326S | 1:50 (0.672 µg) |
| **H3K27ac** | abcam | ab4729 | 1:200 |
| **H3K27me3** | Activ Motif | 39155 | 1:50 (4 µg) |
| **H3K9me3** | abcam | ab8898 | 1:50 |
| **panH3** | abcam | ab1791 | 1:50 (4 µg) |
| **Isotype IgG** | Thermo | 10500C | 1:50 (12 µg) |

**Table S4.** *In vitro* methylation scheme used for training and validation of the methyl-caller model for mA. Different combinations of adenine (mA) and cytosine (mC) methylation were generated using EcoGII, M.CviPI, and M.SssI enzymes. Reaction conditions, incubation times, and subsequent DNA processing steps are indicated for each sample condition prior to Oxford Nanopore Technologies (ONT) library preparation and sequencing.

| Sample | 0<br>Unmeth | 3<br>mA only | 4<br>mC only | 7<br>mC + mA |
| --- | --- | --- | --- | --- |
| Step 1 | — | 25 U/μg EcoGII (mA) | 8 U/μg M.CviPI (mGpC) | 8 U/μg M.CviPI (mGpC) |
|  |  | 1.2 mM SAM<br>30 min at 37°C | 1.2 mM SAM<br>30 min at 37°C | 25 U/μg EcoGII (mA)<br>1.2 mM SAM<br>30 min at 37°C |
| Step 2 | — | 25 U/μg EcoGII (mA) | 8 U/μg M.CviPI (mGpC) | 8 U/μg M.CviPI (mGpC) |
|  |  | 1.2 mM SAM<br>30 min at 37°C | 1.2 mM SAM<br>30 min at 37°C | 25 U/μg EcoGII (mA)<br>1.2 mM SAM<br>30 min at 37°C |
| Step 3 | — | Stop solution<br>14 μM Proteinase K<br>> 4 hours at 55°C | 16 U/μg M.SssI (mCpG)<br>7.5 μM MgCl2<br>0.8 mM SAM<br>30 min at 37°C | 16 U/μg M.SssI (mCpG)<br>7.5 μM MgCl2<br>0.8 mM SAM<br>30 min at 37°C |
| Step 4 | — | — | Stop solution<br>14 μM Proteinase K<br>> 4 hours at 55°C |  |
| Step 5 | — | DNA purification |  |  |
| Step 6 | ONT Library preparation (SQK-NBD114.24) + sequencing |  |  |  |

